# Antiviral signalling in human IPSC-derived neurons recapitulates neurodevelopmental disorder phenotypes

**DOI:** 10.1101/789321

**Authors:** Katherine Warre-Cornish, Leo Perfect, Roland Nagy, Matthew J. Reid, Annett Mueller, Amanda L. Evans, Cédric Ghevaert, Grainne McAlonan, Eva Loth, Declan Murphy, Deepak P. Srivastava, Jack Price

**Author notes:** These authors contributed equally to this work (Katherine Warre-Cornish, Leo Perfect). joint Senior and corresponding authors.

## Abstract

Maternal immune activation increases the risk of neurodevelopmental disorders. Elevated cytokines, such as interferon-gamma (IFNγ), in offspring’s brains play a central role. IFNγ activates an antiviral cellular state, limiting viral entry and replication. In addition, IFNγ has been implicated in brain development. Here, we hypothesise that IFNγ-induced antiviral signalling contributes to molecular and cellular phenotypes associated with neurodevelopmental disorders. We find that transient IFNγ treatment of neural progenitors derived from human induced pluripotent stem cells (hIPSCs) persistently increases neurite outgrowth, phenocopying hIPSC-neurons from autistic individuals. IFNγ upregulates antiviral PML bodies and MHC class I (MHCI) genes, which persists through neuronal differentiation. Critically, IFNγ-induced neurite outgrowth requires both PML and MHCI. We also find that IFNγ disproportionately alters expression of autism and schizophrenia risk genes, suggesting convergence between these genetic and environmental risk factors. Together, these data indicate that IFNγ-induced antiviral signalling may contribute to neurodevelopmental disorder aetiology.

## Introduction

Multiple lines of evidence point to immune activation during foetal development as an important risk factor for neurodevelopmental disorders *(1)*. Epidemiological studies indicate that maternal infection during pregnancy increases incidence of autism spectrum disorder (ASD) and schizophrenia (SZ) *(2–5)*. Animal studies have shown that induction of an antiviral immune response during pregnancy using the dsRNA mimetic poly(I:C) leads to behavioural abnormalities in offspring that are thought to be relevant to neuropsychiatric disorders, including repetitive behaviour, altered social behaviour, deficits in prepulse inhibition and working memory *(6)*. Also, transcriptomic studies of both ASD and SZ post mortem brains consistently show enrichment for inflammatory and innate immune genes *(7–10)*. However, despite this association, the pathological mechanisms through which transient inflammatory activation increases susceptibility to neurodevelopmental disorders remain unclear.

Among the inflammatory cytokines upregulated during maternal immune activation *(11)*, IFNγ is of particular interest. It is an activator of innate cellular antiviral signalling and transcription programmes whose primary function is to defend the cell against infection by viral particles *(12)*. Mid-pregnancy maternal serum IFNγ is increased during gestation of offspring with ASD *(13)* and circulatory IFNγ levels are elevated in neonates subsequently diagnosed with ASD relative to developmental delay controls *(14)*. Intriguingly, within the brain, many antiviral IFNγ signalling targets also play important roles in neuronal development and synaptic activity, independent of microbial infection *(15–17)*. More recently, IFNγ has also been described to play a role in social behaviour in rodents, through modulation of inhibitory neuronal GABAergic tone *(18)*. Thus, it is now emerging that IFNγ has a physiological role beyond its antiviral and immune actions.

Previous studies have shown that antiviral activation establishes enduring cellular changes that persist beyond the acute inflammatory response. Exposure to IFNγ primes cells to induce an enhanced transcriptional response upon re-stimulation, allowing cells to mount a faster and more effective antiviral response *(19, 20)*. Examination of transcriptional priming at the MHC locus revealed a critical role for antiviral promyelocytic leukemia protein (PML) nuclear bodies *(20)*. MHC Class I (MHCI) gene expression has also been shown to be persistently upregulated in developing neurons following gestational poly(I:C) exposure *(21)*. In the brain, seemingly independently of their antiviral functions, both PML and MHCI proteins play important roles in many aspects of neuronal development and function, including neurite outgrowth and axon specification *(15)*, synaptic specificity *(16)*, synaptic plasticity *(22)* and cortical lamination *(23)*. Importantly, all of these processes have been shown to be altered following gestational poly(I:C) exposure in rodents *(6, 11, 21, 24)*. Thus, PML nuclear bodies and MHCI proteins may link antiviral inflammatory activation to neuronal abnormalities. However, the impact of this pathway on neurodevelopment following inflammatory activation has never been examined.

In this study, we examine the hypothesis that IFNγ-induced antiviral signalling perturbs neurodevelopmental processes associated with neurodevelopmental disorders. Here, we used hIPSCs to investigate how transient IFNγ exposure affects developing neurons. We hypothesised that transient developmental inflammatory activation could recapitulate molecular and cellular phenotypes associated with neurodevelopmental disorders. Indeed, we demonstrate that exposing hIPSC-derived neural progenitor cells (NPCs) to IFNγ led to persistently increased neurite outgrowth in hIPSC-neurons, mimicking a phenotype observed in hIPSC-neurons from individuals with ASD *(25–27)*. RNA sequencing was used to characterise the acute and persistent transcriptomic responses to IFNγ. Importantly, we observed that genes of the MHCI protein complex were among the most significantly upregulated. This was accompanied by a persistent increased expression of MHCI proteins and number of PML bodies. Critically, both PML and MHCI proteins were required for IFNγ-dependent effects on neuronal morphology. Furthermore, we observed higher numbers of PML bodies in NPCs derived from individuals diagnosed with ASD than controls, consistent with transcriptomic perturbations in post-mortem human brain *(10)*. Interestingly, IFNγ responding genes were enriched for those with genetic association to ASD and SZ. Moreover, these genes overlapped significantly with those differentially expressed in the brains of individuals with these disorders. Together, these findings highlight a potential mechanism through which antiviral signalling could contribute to intrinsic neuronal phenotypes in neurodevelopmental disorders.

## Results

### IFNγ increases neurite outgrowth in a hIPSC model of neurodevelopment

We and others have previously observed alterations in the morphology of hIPSC neurons derived from autistic individuals *(25–27)*. Since inflammatory mechanisms have been implicated in both neural development and neurodevelopmental pathology, we hypothesised that activation of antiviral signalling pathways may influence neuronal architecture. To address this, we used hIPSC-NPCs from three control male individuals with no psychiatric diagnoses (M1, M2 and M3; Supplementary Table 1; Fig. S1; *21, 23)* and treated these cells with IFNγ (25 ng/ml) daily on days (D) 17-21 of differentiation. Subsequently, IFNγ was excluded from cell culture media and neuronal differentiation was continued, resulting in postmitotic hIPSC-neurons (Fig.1A). Cells were fixed on D26, D30, D35 and D40, stained for βIII-tubulin (Tuj1) and high-content automated neurite tracing was carried out (Fig.1, B and C). Both IFNγ treatment and days in culture were associated with a significant increase in total neurite length per cell across the time-course examined (Fig. 1D; IFNγ treatment *P* = 0.0004; days in culture *P* < 0.0001). We used Sidak’s multiple comparison test to compare IFNγ-treated and untreated neurons at individual time-points and observed a significant increase in total neurite length in IFNγ-treated lines at D30, (*P* = 0.010) after which increased variability was observed. This is similar to the increased neurite length observed in ASD-derived neurons in previous studies *(25–27)*.

**Figure 1:**
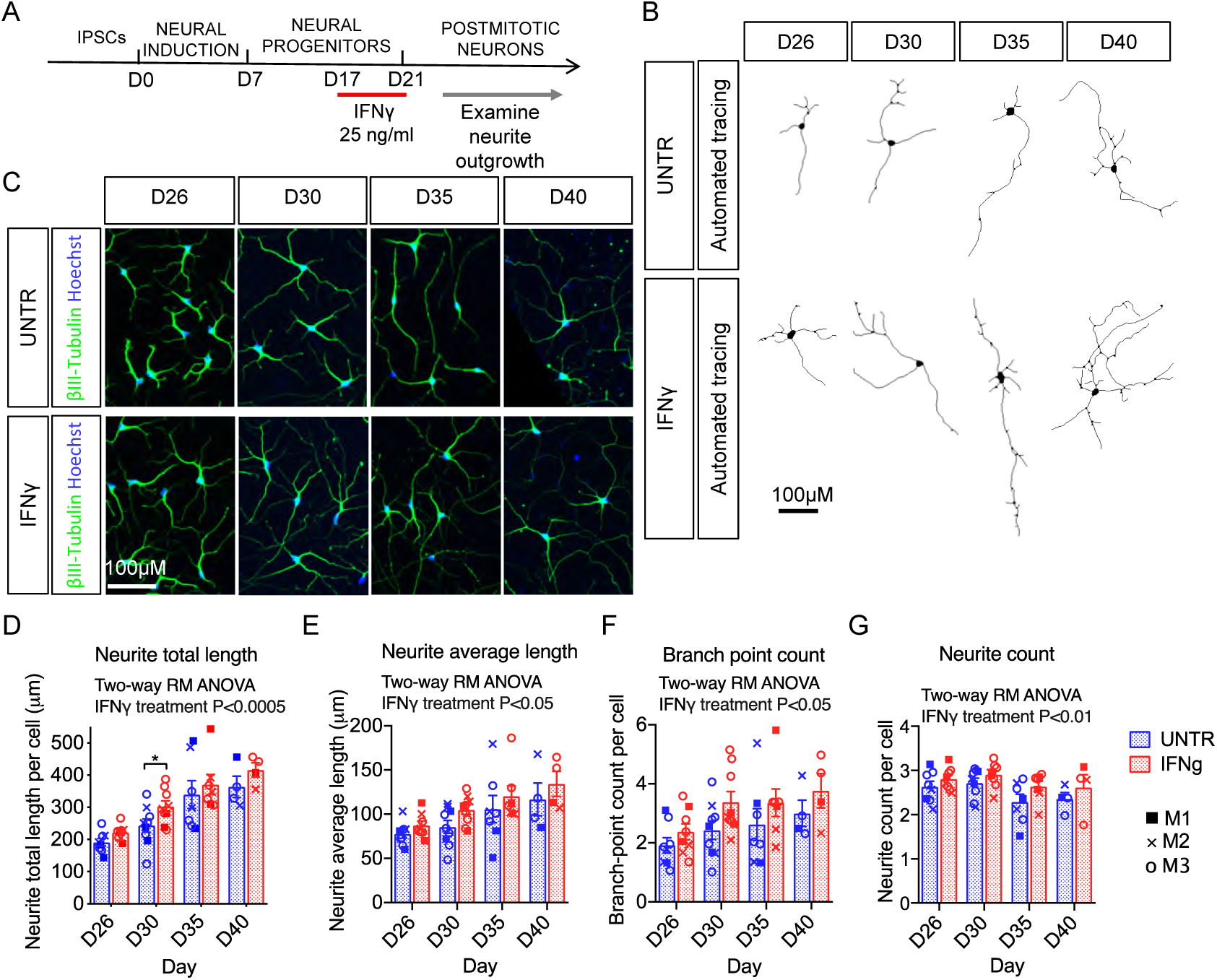
IFNγ treatment of NPCs leads to enhanced neurite outgrowth in postmitotic neurons. **(A)** Schematic representation of the experimental timeline of IPSC differentiation and IFNγ treatment strategy. Neural progenitor cells received 25ng/ml IFNγ daily in cell culture media from D17-D20 before terminal plating on D21 and examination of neurite outgrowth. **(B)** Automated tracing of βIII-Tubulin stained neurites on D26, D30, D35 and D40, carried out with CellInsight™ high content screening operated by HCS Studio Software. **(C)** Fluorescence images of βIII-Tubulin and Hoechst staining acquired with CellInsight™. **(D-G)** Graphs show the time-courses of neuronal morphological properties including: Neurite total length per cell **(D)**; Neurite average length per cell **(E)**; Branch point count per cell **(F)**; Neurite count per cell **(G)** in 3 control male cell lines: M1, M2 and M3. D26 Untreated: *n* = 8 independent biological replicates; 6,382 cells analysed; D26 IFNγ: *n* = 8 independent biological replicates; 7,122 cells analysed; D30 Untreated: *n* = 9 independent biological replicates; 5,651 cells analysed; D30 IFNγ: *n* = 9 independent biological replicates; 7,741 cells analysed; D35 Untreated: *n* = 7 independent biological replicates; 4,250 cells analysed; D35 IFNγ: *n* = 7 independent biological replicates; 4,733 cells analysed; D40 Untreated: *n* = 4 independent biological replicates; 2,792 cells analysed; D40 IFNγ: *n* = 4 independent biological replicates; 2,872 cells analysed. Data generated with CellInsight™ high content screening operated by HCS Studio Software. Results are presented as means +/- SEM. Two-way repeated measures ANOVA with Sidak’s multiple comparison test. \**P* < 0.05.

In previous studies of cells from individuals with ASD, increased neurite branching, number, and length has also been reported *(25–27)*. Consequently, we looked to see if these features were altered in IFNγ-treated cells. IFNγ treatment led to a significant increase in average neurite length per cell across the time-course (Fig. 1E; IFNγ treatment *P* = 0.034; days in culture *P* = 0.0027), although multiple comparison testing showed no significant difference between IFNγ and untreated cells at individual time-points. Branch-point count per cell was also higher in IFNγ-treated cells across the time-course (Fig. 1F; IFNγ treatment *P* = 0.013; days in culture *P* = 0.067) although, similarly, no significant difference was observed with treatment at individual time-points. The neurite count per cell was also higher among IFNγ-treated neurons but, unexpectedly, tended to decrease across the time-course in both conditions (Fig. 1G; IFNγ treatment *P* = 0.0099; days in culture *P* = 0.068). Again, we observed no significant difference in neurite count per cell between IFNγ and untreated cells at individual time-points. Taken together, these data indicate that exposing hIPSC-NPCs to IFNγ results in a persistent increase in total neurite outgrowth resulting from longer, more numerous neurites and increased branching in hIPSC-derived neurons compared to untreated controls, consistent with previous findings with autism-derived cells.

### Transcriptomic analysis of IFNγ exposure during neuronal differentiation

We carried out RNA-sequencing analysis of the three control cell lines (M1, M2 and M3; Supplementary Table 1) to examine the transcriptomic changes associated with IFNγ treatment of human NPCs and neurons. This experiment had six conditions and is schematised in Figure 2A. We sought to capture: (a) the acute response of NPCs to IFNγ, (b) the acute response of neurons to IFNγ, (c) the persistent response of neurons after IFNγ exposure at the NPC stage, and (d) the effect of repeated IFNγ exposure at the NPC and neuronal stages. Principal component analysis revealed that the largest source of variation across the entire dataset corresponded to cell type, with the first principal component explaining 72% of variance and separating NPCs and neurons (Fig. 2B). The second principal component segregated the samples by IFNγ exposure and described 20% of the observed variance between samples.

**Figure 2:**
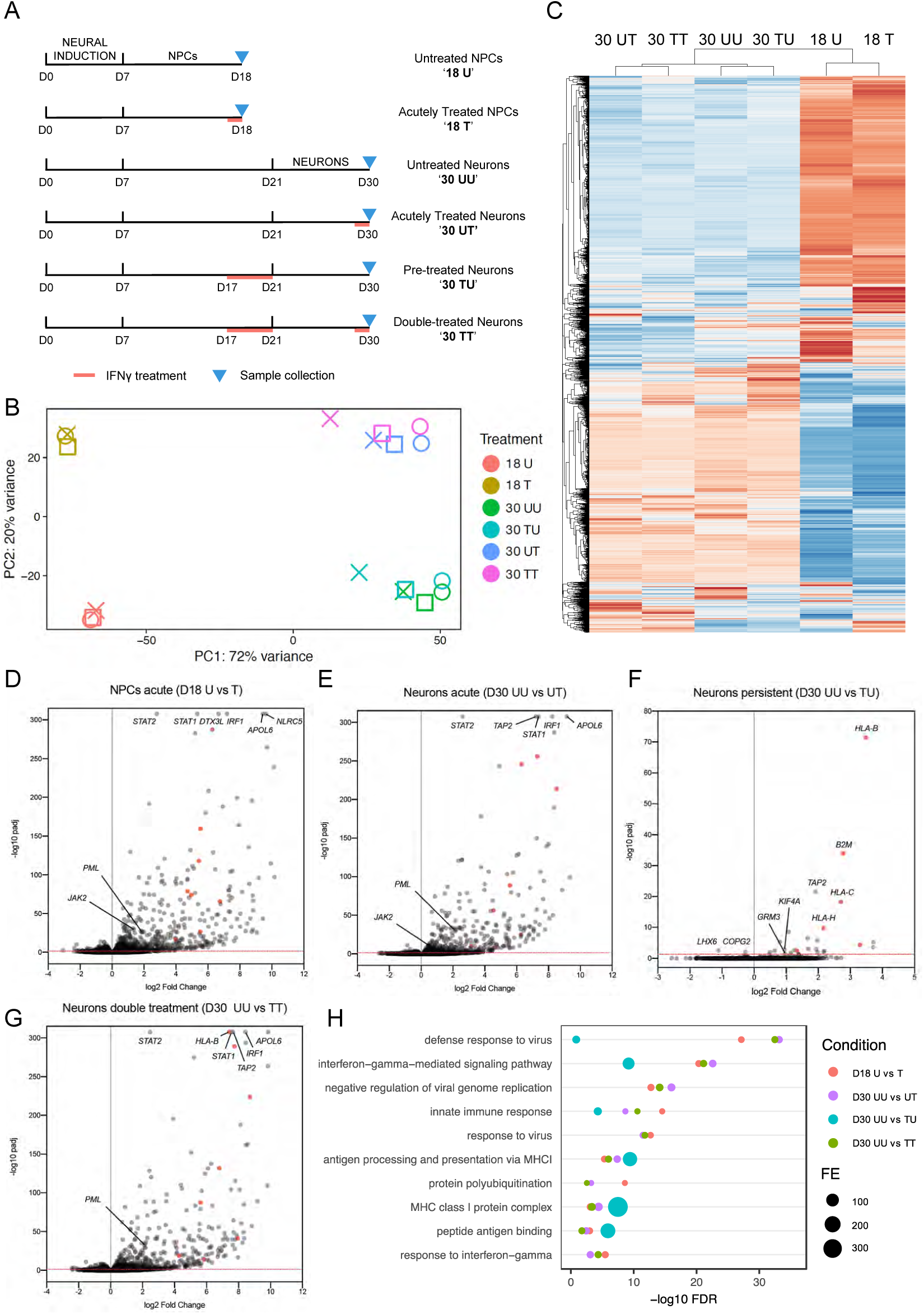
RNA-sequencing analysis reveals a widespread and persistent transcriptomic response of human NPCs and neurons to IFNγ. **(A)** Schematic representation of the experimental conditions. **(B)** PCA biplot of all samples. The first principal component segregates conditions by time-point, while the second separates conditions by recent treatment. Cell lines are represented by point shape: M1 = square, M2 = cross, M3 = circle. **(C)** Heatmap of all differentially expressed genes clustered by row and column. Replicates are collapsed by condition. **(D-G)** Volcano plots with selected genes annotated. The dotted red line represents the threshold for statistical significance of padj = 0.05. Red dots identify genes of the MHCI protein complex. **(H)** Cleveland plot of selected enriched GO terms from the upregulated gene sets illustrating statistical significance and fold enrichment (FE).

Comparing untreated (18U) and acutely treated (18T) NPCs, we discovered 1,834 genes were differentially expressed in response to IFNγ (7.7% of 23,757 genes measured; Fig. 2, C and D). The comparison of untreated (30UU) and acutely treated neurons (30UT) revealed 751 differentially expressed genes (DEGs; 2.8% of 27,082 genes measured; Fig. 2, C and E). The canonical IFNγ signalling pathway was activated with acute treatment in both NPCs and neurons, resulting in significant upregulation of *JAK2, STAT1, STAT2* and downstream interferon stimulated genes such as *IRF1* (Fig. 2, D and E). Consequently, upregulated genes were highly enriched for the gene ontology (GO) term ‘interferon-gamma-mediated signalling pathway’ (9-fold in NPCs and 14-fold in neurons; FDR = 4.47 x 10^−21^ and FDR = 2.68 x 10^−23^ respectively; Fig. 2H). Genes belonging to the MHCI complex (GO term ‘MHC class I protein complex’) were overrepresented in the upregulated genes (11-fold in NPCs and 21-fold in neurons; FDR = 8 x 10^−4^ and FDR = 3.98 x 10^−5^ respectively; Fig. 2H) and among the highest ranked DEGs in NPC and neurons acutely treated with IFNγ (Fig. 2, D and E). We also found that the genes downregulated by acutely treated neurons were enriched 2-fold for the GO term ‘plasma membrane’ (FDR = 0.002). Full differential expression and GO enrichment results are reported in supplementary files S1 and S2 respectively.

The persistent transcriptional response to IFNγ was of interest, given the enduring impact of IFNγ on neuronal morphology described above. To this end, we compared untreated neurons (30UU) and pre-treated neurons (30TU; Fig. 2A). Remarkably, neurons exposed to IFNγ at the NPC stage showed enduring transcriptional changes, with 26 genes significantly upregulated and 2 downregulated in postmitotic neurons 9 days after treatment (Fig. 2F). Notably, the upregulated gene set was enriched for MHCI genes, with the GO term ‘MHC class I protein complex’ enriched 357-fold (FDR = 3.33 x 10^−8^, Fig. 2H). Strikingly, the top five ranked DEGs in pre-treated neurons were all involved in MHCI antigen presentation. Also of interest, the metabotropic glutamate receptor gene *GRM3* was upregulated while the GABAergic transcription factor gene *LHX6* was downregulated in the pre-treated neurons. Interestingly, *GRM3* has been implicated as a SZ risk gene *(29)*, whereas *LHX6* is required for specification and correct spatial positioning of parvalbumin interneurons, neurite outgrowth, and has been reported to be deficient in prefrontal cortices of individuals with SZ *(30)* (Fig 2F).

We also found evidence of IFNγ-induced cellular priming, where repeated exposure at the NPC and neuronal stages (30TT) induced considerably more DEGs than a single neuronal treatment (30UT) when compared to untreated neurons (30UU). This double hit induced 1,091 DEGs, 45% more than were detected after a single neuronal treatment (Fig. 2G). This is consistent with previous reports of IFNγ-induced transcriptional priming *(19, 20)*. Notably, the genes downregulated by neurons that received a double hit were enriched 4-fold for the GO term ‘synapse’ (FDR = 0.049). Taken together, these results demonstrate that IFNγ exposure induces widespread and persistent transcriptional changes during human neuronal differentiation. Considering their previously proposed role in neurite outgrowth in mouse *(15, 31)*, these data highlight MHCI proteins as candidates to explain the IFNγ-induced morphological phenotype described above.

### Regulation of MHCI genes by PML nuclear bodies

In non-neuronal cells, IFNγ has been shown to induce long-lasting changes in signal-dependent transcription of MHC proteins through the formation of PML bodies *(20)*. PML bodies are dynamic, DNA-binding protein complexes which mediate myriad transcriptional functions from viral gene silencing *(32)* to neurogenesis *(23)* and neuronal homeostatic plasticity *(22)*. Because the MHCI pathway was highly enriched in the pre-treated neurons and *PML* was similarly upregulated in acutely treated NPCs and neurons (Fig. 2D and E), we investigated the putative role of PML bodies in mediating the effects of IFNγ. We hypothesised that PML bodies could underlie the observed IFNγ-induced MHCI expression. Cells from the six IFNγ treatment conditions (Fig. 2A) were stained for PML (Fig. 3, A and C). Acute 24-hour treatment of NPCs at D17-18 resulted in a highly significant increase in PML bodies per cell (Fig. 3, A and B; *P* < 0.0001). Importantly, the increase in PML bodies persisted through differentiation and was still observed in neurons (Fig. 3, C and D; *P* < 0.0001). Conversely, treatment of postmitotic neurons had no effect on PML bodies, with or without pre-treatment at D17-21 (Fig. 3D). These results indicate a time-window of sensitivity in neuronal differentiation, during which IFNγ exposure leads to a persistent increase in PML bodies.

**Figure 3:**
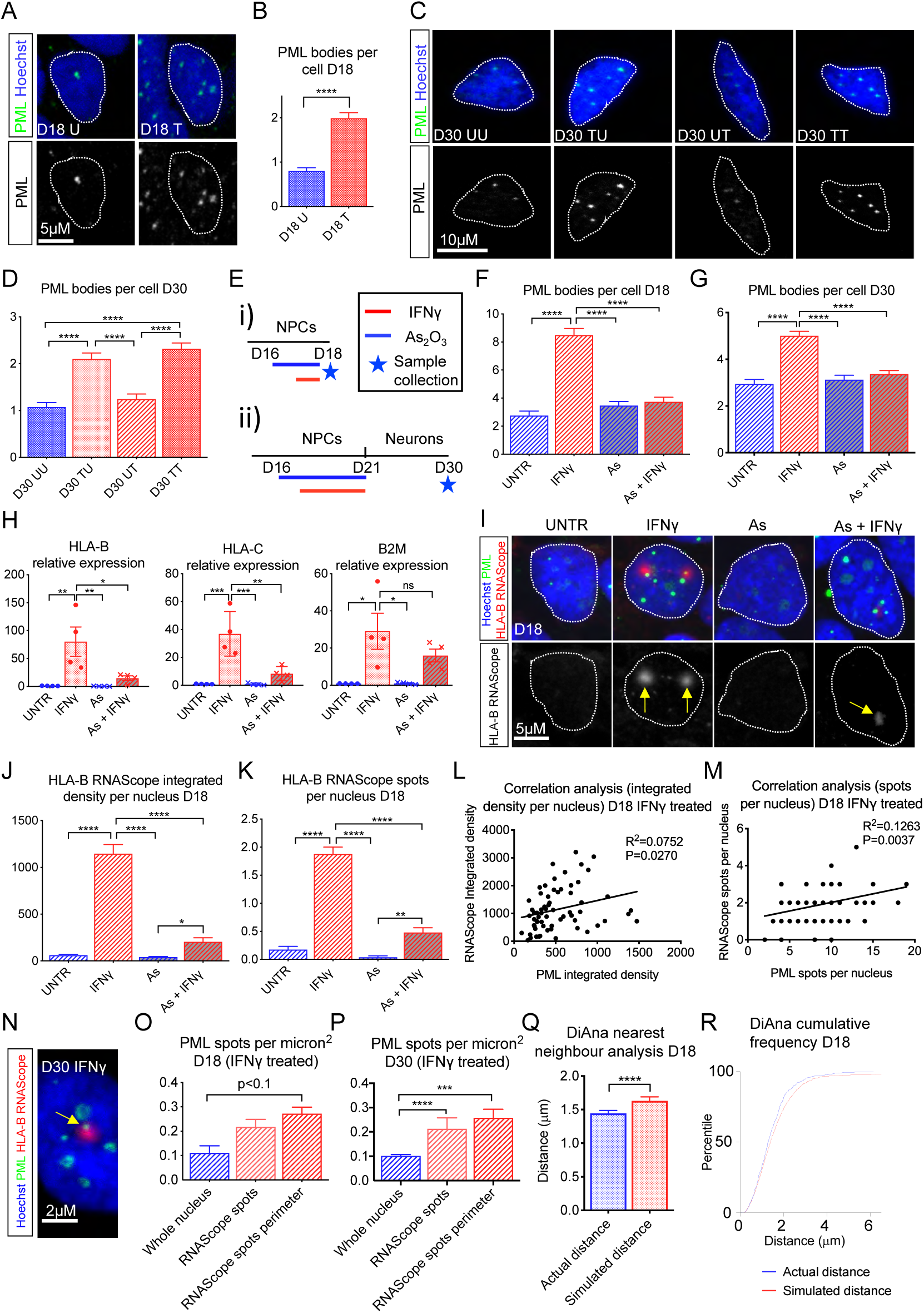
PML bodies are persistently increased following treatment of NPCs with IFNγ, regulate transcription of MHCI genes and are spatially associated with *HLA-B* transcription. **(A)** Confocal images of PML nuclear bodies in untreated (D18 U) and IFNγ-treated (D18 T) NPCs. **(B)** Quantification of PML bodies per nucleus in D18 NPCs. D18 U: *n* = 354 cells; D18 T: *n* = 286 cells; 3 control cell lines. Two-tailed Mann-Whitney test. **(C)** Confocal images of nuclear PML bodies in untreated (D30 UU), pre-treated (D30 TU), acutely treated (D30 UT), and double-treated (D30 TT) D30 neurons. **(D)** Quantification of nuclear PML bodies in D30 neurons. D30 UU: *n* = 236 cells; D30 TU: *n* = 264 cells; D30 UT: *n* = 231 cells; D30 TT: *n* = 307 cells; 3 control cell lines. **(E)** Schematic representation of IFNγ and As_2_O_3_ treatment conditions. Kruskal-Wallis test with Dunn’s multiple comparison test. **(F and G)** Quantification of PML bodies in D18 and D30 ‘UNTR’, ‘IFNγ’, ‘As’ and ‘As + IFNγ’ conditions. D18 UNTR: *n* = 57 cells; D18 IFNγ: *n* = 65 cells; D18 As: *n* = 64 cells; D18 IFNγ + As: *n* = 54 cells. D30 UNTR: *n* = 104 cells; D30 IFNγ: *n* = 129 cells; D30 As: *n* =119 cells; D30 As + IFNγ: *n* = 105 cells; 3 control cell lines. Kruskal-Wallis test with Dunn’s multiple comparison test. **(H)** qPCR analysis of *HLA-B, HLA-C* and *B2M* relative expression in D18 ‘UNTR’, ‘IFNγ’, ‘As’ and ‘As + IFNγ’ conditions. *n* = 4 biological replicates from 3 control cell lines. One-way ANOVA with Tukey’s multiple comparison test. **(I)** Confocal images of *HLA-B* RNAScope™ F.I.S.H. and PML immunocytochemistry in D18 ‘UNTR’, ‘IFNγ’, ‘As’ and ‘As + IFNγ’ conditions. **(J and K)** Quantification of *HLA-B* RNAScope™ integrated density and spots per nucleus in D18 ‘UNTR’, ‘IFNγ’, ‘As’ and ‘As + IFNγ’ conditions. D18 UNTR: *n* = 57 cells; D18 IFNγ: *n* = 65 cells; D18 As: *n* = 64 cells; D18 IFNγ + As: *n* = 54 cells. Kruskal-Wallis test with Dunn’s multiple comparison test. **(L and M)** Correlation analysis of *HLA-B* RNAScope and PML integrated density and spots per nucleus in D18 IFNγ-treated cells. *n* = 65 cells from 3 control cell lines. Linear regression analysis. **(N)** Confocal image of *HLA-B* RNAScope™ F.I.S.H. and PML immunocytochemistry in D30 IFNγ treatment condition. Arrow shows colocalisation between PML and *HLA-B* pre-mRNA. **(O and P)** Quantification of PML spots per micron^2^ in whole nuclei, *HLA-B* RNAScope spots and *HLA-B* RNAScope spot perimeters in D18 NPCs and D30 neurons, IFNγ treatment condition. D18 IFNγ: *n* = 67 cells; D30 IFNγ: *n =* 58 cells; 3 control cell lines. Kruskal-Wallis test with Dunn’s multiple comparison test. **(Q and R)** DiAna distance analysis of centre-centre distances from *HLA-B* RNAScope spots to PML spots in real images and following random shuffle of PML and RNAScope spots, using nuclei as a bounding box. n = 405 measurements; 3 control cell lines. Wilcoxon matched-pairs test. Results are presented as means +/- SEM. \**P* < 0.05; \*\**P* < 0.01; \*\*\**P* < 0.001; \*\*\*\**P* < 0.0001; ns: not significant.

PML nuclear bodies are known to be specifically disrupted following binding of arsenic trioxide (As_2_O_3_) *(33)*. Therefore, we pre-treated NPCs with As_2_O_3_ from D16-17, followed by co-treatment with As_2_O_3_ and IFNγ from D17-18, then counted PML bodies per nucleus (Fig. 3, E i, F). As previously observed (Fig. 3A), IFNγ treatment led to a notable increase in PML bodies per nucleus (*P* < 0.0001); this was blocked by co-treatment with As_2_O_3_. Alone, As_2_O_3_ had no detectable impact on number of PML bodies. To test whether As_2_O_3_ treatment also prevented the persistent increase in PML bodies the experiment was repeated, this time allowing NPCs to differentiate into neurons (Fig. 3, E ii, G). Again, we observed a significant increase in PML bodies with IFNγ treatment alone (*P* < 0.0001) which was entirely prevented by co-treatment with As_2_O_3_. We could thus conclude that As_2_O_3_ prevented both acute and persistent PML body induction.

To investigate whether PML bodies were required for IFNγ-dependent transcriptional activation of MHCI genes, we performed quantitative (q)PCR on NPCs exposed to As_2_O_3_, IFNγ, or combined treatment (Fig. 3, E i). We observed that IFNγ-dependent induction of the MHCI genes *HLA-B* and *HLA-C* was blocked by exposure to As_2_O_3_ (Fig. 3H). Interestingly, induction of the MHCI receptor subunit *B2M*, was not blocked by As_2_O_3_, suggesting that the PML-dependent effect is specific to MHC genes. Similarly, neither of the genes encoding IFNγ receptor subunits, IFNGR1 and IFNGR2, were affected by As_2_O_3_ co-treatment (Fig. S2, A and B). Collectively, these data support a model in which PML nuclear bodies mediate IFNγ-dependent gene transcription of MHCI genes.

To further interrogate the necessity of PML for IFNγ-induced MHCI gene expression, RNA FISH was carried out on NPCs treated with IFNγ, As_2_O_3_, or co-treated with IFNγ and As_2_O_3_ (Fig. 3, E i, I). We used a probe specific to the pre-spliced *HLA-B* RNA transcript. *HLA-B* was selected as it showed the greatest upregulation in pre-treated neurons (Fig. 2F). IFNγ treatment led to a dramatic increase in *HLA-B* pre-mRNA (Fig. 3, I, J and K; *P* < 0.0001). This induction was largely prevented by co-treatment with As_2_O_3_ (Fig. 3I, J and K), supporting the requirement for PML in IFNγ-dependent *HLA-B* transcription. Furthermore, co-staining for PML and *HLA-B* pre-mRNA revealed a positive correlation in the IFNγ-treated condition. This was observed as both integrated density of PML and *HLA-B* per nucleus (Fig. 3L; Slope significantly greater than zero, *P* = 0.027) or as number of spots per nucleus in each channel (Fig. 3M; Slope significantly greater than zero, *P* = 0.0037). A significant positive correlation between PML and *HLA-B* spots per nucleus was also observed in the unstimulated condition (Fig. S3A; *P* = 0.0014), although there was no correlation between the integrated density of PML and *HLA-B* (Fig. S3B; *P* = 0.38). A relationship between PML bodies and IFNγ-induced *HLA-B* transcription is thus strongly supported.

### IFNγ induces *HLA-B* transcription near PML nuclear bodies

If PML bodies directly regulate MHCI gene expression, we would predict them to be in close proximity to the site of transcription. To investigate this spatial relationship, we carried out RNA FISH using the probe described above, specific to pre-spliced *HLA-B* RNA transcript. Because splicing is highly localised *(34, 35)*, we reasoned that the site of the pre-spliced transcript could be used as a proxy for the location of transcription. Co-staining for PML and *HLA-B* pre-mRNA revealed that, following IFNγ treatment, *HLA-B* spots were frequently located immediately adjacent to, or overlapping with, PML bodies (Fig. 3N). To investigate this further, we measured the density of PML bodies (spots per micron) within *HLA-B* spots or *HLA-B* spot perimeters (see Methods for full definitions) and compared this to the density of PML bodies across the nucleus as a whole in IFNγ-treated NPCs and neurons (Fig. 3, O and P; S3C and S3D). In IFNγ-treated NPCs, the increased density of PML bodies in *HLA-B* pre-mRNA spot perimeters did not reach statistical significance (*P* = 0.09). However, in IFNγ-treated neurons, a significantly higher density of PML bodies was observed in the *HLA-B* pre-mRNA spots (*P* < 0.0001) and spot perimeters (*P* = 0.0007) than the nucleus as a whole, indicating a positive spatial association. By contrast, in untreated NPCs and neurons, PML spots were never observed to overlap with *HLA-B* pre-mRNA spots or spot perimeters (Fig. S3C and S3D). To confirm the existence of a non-random spatial relationship, we carried out random shuffle nearest neighbour analysis on IFNγ-treated NPCs *(36)*. Actual distances between *HLA-B* spots and PML bodies were observed to be significantly shorter than simulated distances following randomisation (*P* < 0.0001), confirming a positive spatial relationship (Fig. 3, Q and R). These results confirm that PML bodies are in closer proximity to the site of *HLA-B* transcription than would be expected by chance, supporting the hypothesis that PML is required for IFNγ-induced MHCI gene transcription.

### PML and MHCI mediate IFNγ-induced neurite outgrowth

Having demonstrated a role for PML bodies in IFNγ-induced MHCI gene transcription, we next asked whether PML and MHCI were required for the IFNγ-dependent increased neurite outgrowth. MHCI proteins have previously been reported to be expressed during, and required for, neurite outgrowth in primary cultured rodent neurons *(15)*. To examine the role of MHCI proteins in IFNγ-induced neurite outgrowth, we first carried out instant (i)SIM super-resolution microscopy to determine the subcellular localisation of MHCI proteins HLA-A, HLA-B and HLA-C (Fig. 4A). We found MHCI proteins to be present in the neurites, growth cones and cell bodies of untreated neurons. We then examined the effects of IFNγ and As_2_O_3_-induced PML disruption on MHCI abundance in neurites and growth cones, following the experimental outline schematised in Fig. 3E ii. IFNγ treatment induced a significant increase in MHCI intensity within growth cones and neurite compartments (Fig. 4B and C; neurites: *P* = 0.0021; growth cones: *P* = 0.0012). This increase was prevented by co-treatment with As_2_O_3_ in both compartments (*P* = 1 for both). MHCI was enriched in growth cones relative to the associated neurite in both untreated (*P* = 0.0034) and IFNγ-treated neurons (*P* < 0.0001; Fig. 4D). The enrichment of MHCI in growth cones was blocked by As_2_O_3_ treatment, both with and without IFNγ (Fig. 4D; As: *P* = 0.40; As = IFNγ: *P* = 0.13). These results support a requirement for PML-dependent signalling in the expression and appropriate subcellular localisation of MHCI proteins both basally and following IFNγ exposure.

**Figure 4:**
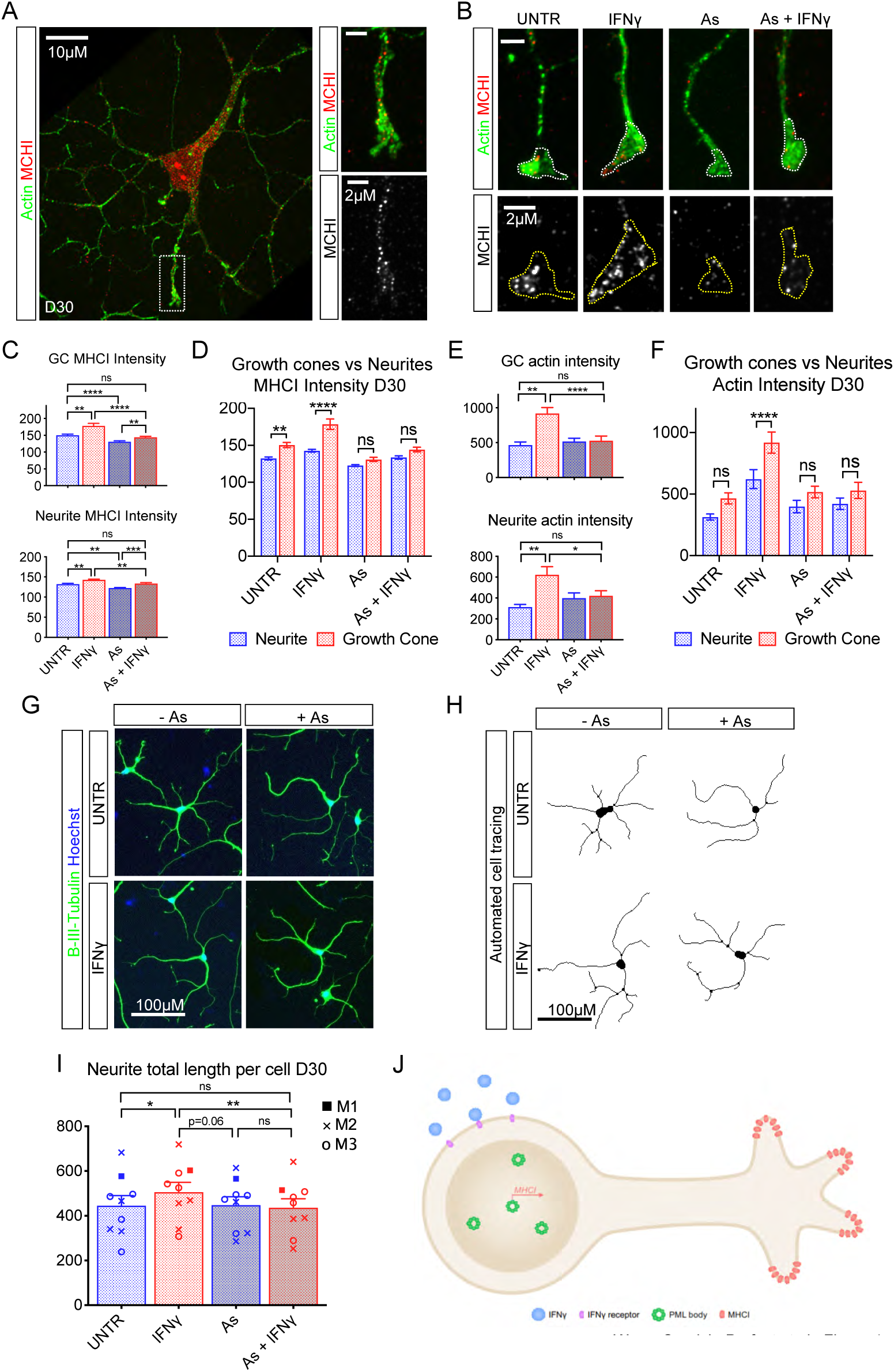
MHCI proteins are enriched in neuronal growth cones in a PML-dependent manner and disruption of PML prevents IFNγ-dependent neurite outgrowth. **(A and B)** iSIM super-resolution images of Actin and MHCI in D30 neurons, with MHCI observable in cell bodies, neurites and growth cones, in UNTR, IFNγ, As and As + IFNγ treatment conditions. **(C-F)** Quantification of MHCI and Actin in D30 neurons, in neurites and growth cones. UNTR: *n* = 52 cells; IFNγ: *n* = 82 cells; As: *n* = 55 cells; As + IFNγ: *n* = 59 cells; 3 control cell lines. **(C and E)** One-way ANOVA with Tukey’s multiple comparison test. **(D and F)** Repeated measures two-way ANOVA with Sidak’s multiple comparison test. **(G)** Fluorescence images of βIII-Tubulin and Hoechst staining acquired with CellInsight™ high content screening system operated by HCS Studio Software. **(H)** Automated tracing of βIII-Tubulin stained neurites in D30 UNTR, IFNγ, As, and As + IFNγ conditions carried out with CellInsight™ high content screening system. **(I)** Graph showing neurite total length per cell in D30 UNTR, IFNγ, As, and As + IFNγ conditions. *n* = 9 independent biological replicates. UNTR: 5,038 cells analysed; IFNγ: 5,668 cells analysed; As: 5,169 cells analysed; As + IFNγ: 3,210 cells analysed; 3 control cell lines. RM one-way ANOVA with Tukey’s multiple comparison test. Data generated with CellInsight™ high content screening operated by HCS Studio Software. **(J)** Schematic depicting our proposed model for IFNγ-induced PML and MHCI-dependent neurite outgrowth. Results are presented as means +/- SEM. \**P* < 0.05; \*\**P* < 0.01; \*\*\**P* < 0.001; \*\*\*\**P* < 0.0001; ns: not significant.

Significantly increased actin intensity was also observed in both neurites and growth cones following IFNγ treatment (Fig. 4E; neurites: *P* = 0.0020; growth cones: *P* = 0.0021). Co-treatment with As_2_O_3_ prevented this increase in both compartments (*P* = 1 for both). Actin staining intensity was also increased in growth cones relative to adjoining neurites following IFNγ treatment (*P* < 0.0001), and this enrichment was prevented by co-treatment with As_2_O_3_ (Fig. 4F; As: *P* = 0.57; As + IFNγ: *P* = 0.13). Together, these results indicate that PML-dependent IFNγ-activated signalling pathways have a functional impact on growth cone composition and actin dynamics.

To determine whether intact PML bodies are required for IFNγ-dependent increased neurite outgrowth, we treated NPCs with IFNγ and As_2_O_3_ then continued differentiation of these cells into post-mitotic neurons and assessed neurite outgrowth. As previously described, we observed an increase in total neurite length per cell in the IFNγ-treated relative to the untreated neurons (Fig. 4, G, H and I; *P* = 0.04). Importantly, this increase was prevented by co-treatment with As_2_O_3_ (*P* = 0.98), supporting a requirement for PML in IFNγ-induced neurite outgrowth. These results support a model whereby IFNγ activates PML body formation, leading to MHCI gene transcription, expression of MHCI in growth cones and increased neurite outgrowth (Fig. 4J).

### IFNγ-mediated increase in MHCI and neurite outgrowth requires expression of B2M

To rule out non-specific effects of As_2_O_3_ and confirm the requirement for MHCI in IFNγ-dependent neurite outgrowth, we investigated the effect of blocking MHCI cell surface expression on IFNγ-induced morphological changes. To achieve this, we took advantage of the requirement for the B2M protein for cell surface expression of MHCI *(37)*. Firstly, we sought to demonstrate that IFNγ-induced expression of B2M protein was required for MHCI expression in our NPCs. Indeed, we observed a significant increase in both B2M and MHCI expression following exposure to IFNγ (Fig. 5, A, B and C; *P* = 0.0011). Critically, shRNA-mediated knockdown of *B2M* blocked IFNγ-induced MHCI expression in NPCs (Fig. 5, A, C, IFNγ vs IFNγ + shRNA B2M k/d: *P* < 0.0001). These data confirm that B2M is required for IFNγ-induced upregulation of MHCI in NPCs. Next, we used a human embryonic stem cell (hESC) line, MS3 HLA null, in which the *B2M* gene has been excised using a CRISPR Cas9 nickase approach. This results in cells lacking cell surface MHCI. We used the MS3 HLA null hESC line to further investigate whether MHCI was required for IFNγ-induced neurite outgrowth. MS3 HLA null hESCs were differentiated using the same neural induction protocol used with our neurons and subsequently, treated with IFNγ (25 ng/ml) on D17-21 (Fig. 1A; S1). Treatment of MS3 HLA null NPCs with IFNγ did not increase MHCI expression (Fig. 5D, E and F; B2M: *P* = 0.68; MHCI: *P* = 0.34), consistent with the requirement for B2M for the expression of MHCI proteins. Critically, when we assessed neurite outgrowth in MS3 HLA null neurons following IFNγ treatment, we no longer observed an effect of treatment on total neurite length (Fig. 5G and H; *P* = 0.74). Taken together, these data demonstrate that IFNγ-mediated alterations in neurite outgrowth require the expression of B2M and MHCI proteins.

**Figure 5:**
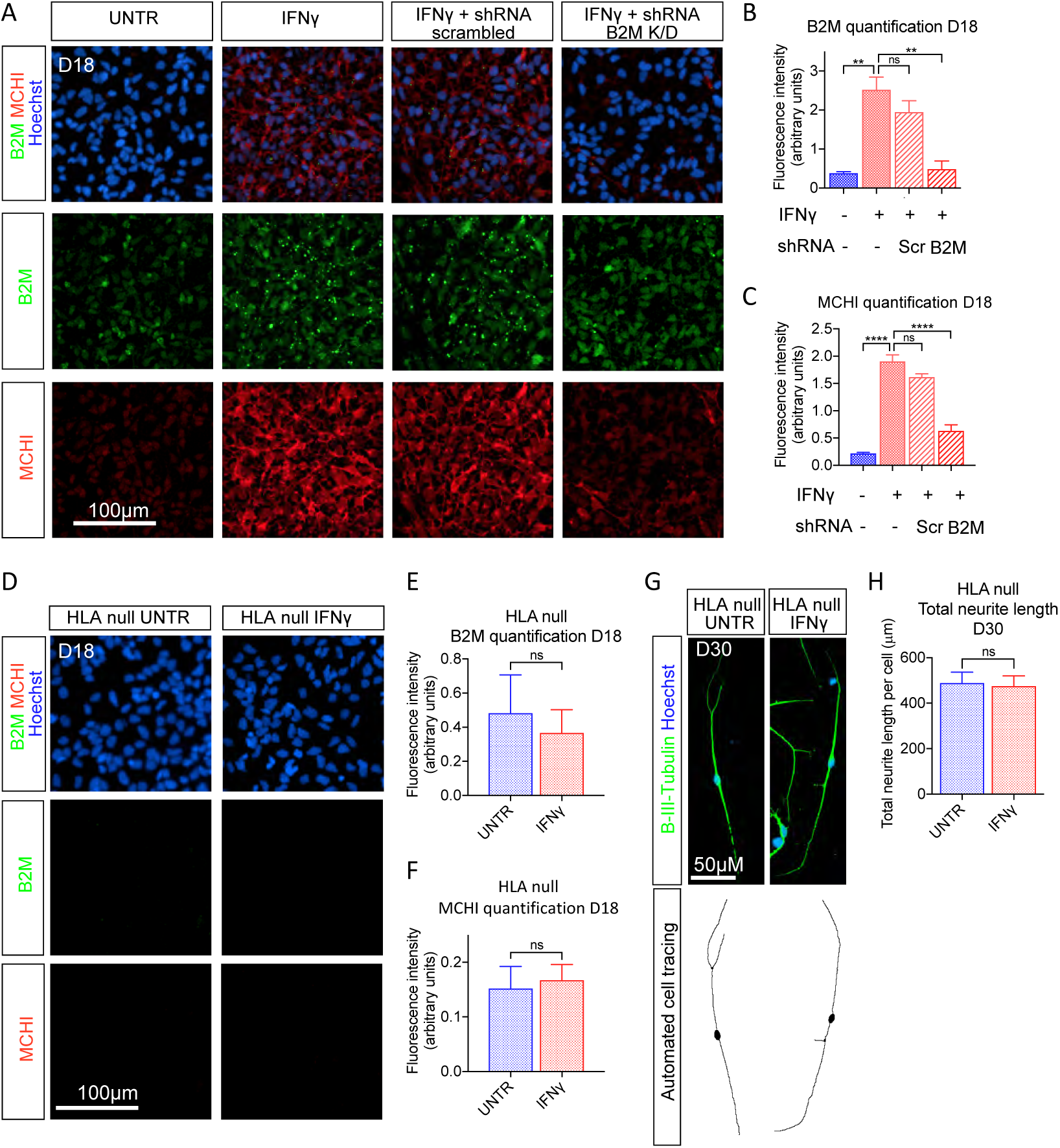
Loss of B2M prevents MHCI cell surface expression and IFNγ-dependent neurite outgrowth. **(A)** Confocal images of B2M, MHCI and Hoechst stained D18 NPCs, UNTR, IFNγ, IFNγ + shRNA scrambled and IFNγ + shRNA B2M knockdown treatment conditions. **(B and C)** Quantification of B2M and MHCI in D18 UNTR, IFNγ, IFNγ + shRNA scrambled and IFNγ + shRNA B2M knockdown. *n* = 3 independent biological replicates; >10,000 cells per condition; 3 control cell lines. One-way ANOVA with Tukey’s multiple comparison correction. **(D)** Confocal images of B2M, MHCI and Hoechst stained D18 MS3 HLA null UNTR and IFNγ-treated NPCs. **(E and F)** Quantification of B2M and MHCI in D18, MS3 HLA null UNTR and IFNγ-treated NPCs. MS3 HLA null UNTR: n = 3 biological replicates; >10,000 cells analysed; MS3 HLA null IFNγ: n = 3 biological replicates; >10,000 cells analysed. Paired t test. **(G)** Fluorescence images of βIII-Tubulin and Hoechst stained neurons and automated tracing of βIII-Tubulin stained neurites in D30 MS3 HLA null UNTR and IFNγ neurons. **(H)** Neurite total length per cell in D30 MS3 HLA null UNTR and IFNγ neurons. *n* = 5 independent biological replicates; MS3 HLA null UNTR >10,000 cells analysed; MS3 HLA null IFNγ >10,000 cells analysed. Paired t test. All images and data generated using Opera Phenix™ high content screening system. Results are presented as means +/- SEM. \**P* < 0.05; \*\**P* < 0.01; \*\*\**P* < 0.001; \*\*\*\**P* < 0.0001; ns: not significant.

### Disease relevance of IFNγ-induced transcriptional alterations

Previous studies have demonstrated that exposure to viral insults during development increases the risk of neurodevelopmental and psychiatric disorders *(1)*. With this in mind, we examined whether genes with genetic association to ASD and SZ were preferentially dysregulated in NPCs and neurons in response to IFNγ (Fig. 6A). Intriguingly, we found ASD-associated genes from the SFARI database *(38)* to be significantly overrepresented among those downregulated by NPCs in response to IFNγ (OR = 1.7, FDR = 0.02). These genes included the candidates *NLGN3, SHANK2, UPF3B* and *NRXN3* (Fig. S4A). Interestingly, we also observed significantly fewer ASD risk genes than were expected by chance among those upregulated by NPCs in response to IFNγ (OR = 0.6, FDR = 0.02). These findings are consistent with a model in which reduced function of ASD risk genes contributes to neurodevelopmental dysregulation following antiviral immune activation. We then tested SZ risk genes curated by the PsychENCODE Consortium (PEC) *(39)*. Again, we found an enrichment among the genes downregulated by NPCs in response to IFNγ (OR = 1.7, FDR = 0.004). Genes in this group included *GRIN2A, PRKD1* and *TSNARE1* (Fig. S5A). The SZ risk genes were also enriched among those upregulated by neurons in response to IFNγ (OR = 1.7, FDR = 0.009). Notable genes in this group included *TCF4, ATXN7* and *ZNF804A* (Fig. S5B). Taken together, these data demonstrate that IFNγ exposure during human neuronal differentiation disproportionately alters genes with genetic association to ASD and SZ. Therefore, we see convergence between genetic and environmental risk factors for these disorders.

**Figure 6:**
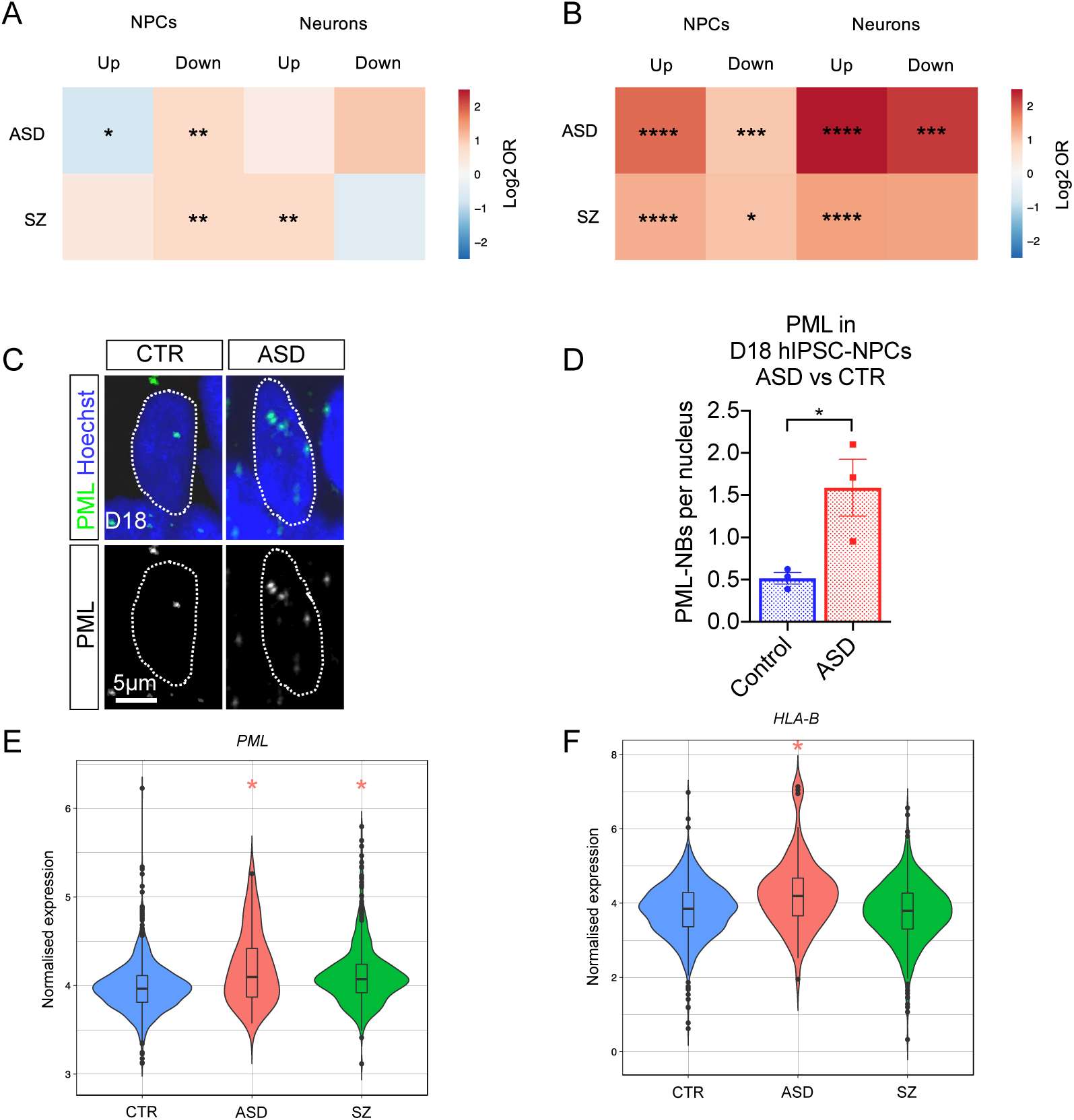
Relevance of IFNγ-dependent gene expression changes to ASD and SZ. **(A)** Enrichment of ASD and SZ risk genes and **(B)** DEGs detected in the brains of ASD and SZ patients among our IFNγ-responding genes. Log2 odds ratio (OR) is represented by colour and statistical significance is indicated by asterisks (*FDR < 0.05; **FDR < 0.01; ***FDR < 0.001; ****FDR < 0.0001). Fischer’s exact test BH corrected for multiple comparisons. **(C)** Confocal images of PML nuclear bodies in D18 NPCs derived from control and ASD individuals. **(D)** Quantification of PML nuclear bodies at D18 in NPCs derived from 3 control and 3 ASD hIPSC lines. CTR: 136 cells analysed; ASD: 280 cells analysed. Unpaired t test. **(E and F)** Violin plots illustrating *PML* and *HLA-B* expression in post-mortem cerebral cortices of controls and individuals with ASD and SZ from the PEC cross-disorder study (15). *n* = 936 control individuals; *n* = 51 ASD individuals; *n* = 559 SZ individuals. Bar chart results are presented as means +/- SEM. \**P* < 0.05; \*\**P* < 0.01; \*\*\**P* < 0.001; \*\*\*\**P* < 0.0001; ns: not significant.

We were also interested to examine the overlap between the IFNγ-responding genes detected in this study with those found to be differentially expressed in the post-mortem brains of patients with ASD and SZ. We utilised data from the PEC cross-disorder study, the largest transcriptome-wide analysis of ASD and SZ brains carried out to date *(10)*. Remarkably, we found a highly significant enrichment of genes from both disorders (Fig. 6B). There was a significant overlap between genes upregulated (OR = 3.7, FDR = 4.5 x 10^−21^) and downregulated (OR = 2.0, FDR = 5.6 x 10^−4^) by NPCs in response to IFNγ with those upregulated and downregulated in the post-mortem brains of individuals with ASD respectively. The effect was even more pronounced for genes upregulated (OR = 5.6, FDR = 2.8 x 10^−25^) and downregulated (OR = 4.6, FDR = 7.6 x 10^−4^) by neurons in response to IFNγ. Next, we tested for overlap with the genes found to be differentially expressed in the post-mortem brains of SZ patients. Again, a significant overlap between genes upregulated (OR = 2.5, FDR = 7.3 x 10^−15^) and downregulated (OR = 2.1, FDR = 0.01) by NPCs in response to IFNγ was observed. Lastly, we also found a significant overlap between genes upregulated by neurons in response to IFNγ with those upregulated in the brains of SZ patients (OR = 2.7, FDR = 9.9 x 10^−12^). The overlapping upregulated genes at both time points were enriched for the gene ontology terms “IFNγ-mediated signalling pathway” and “defence response to virus” in both disorders (File S3). These data indicate that activation of IFNγ-dependent antiviral signalling modifies gene expression in a manner consistent with the dysregulation observed in the brains of patients with ASD and SZ.

### NPCs from idiopathic ASD individuals display increased basal PML nuclear bodies

Finally, to investigate the relevance of PML bodies to neurodevelopmental disorders, we examined hIPSC-derived NPCs originating from individuals diagnosed with ASD. Cells from three individuals diagnosed with ASD alongside the three control cell lines were differentiated into NPCs as described previously (Fig. 1A; S1), fixed and stained for PML on D18 (Fig. 6, C and D). The NPCs from individuals with ASD had more PML bodies per nucleus than controls, indicating over-activation in basal conditions (Fig. 6, C, D; *P* = 0.036). This finding is consistent with published transcriptomic data from post mortem brain tissue *(9, 10, 40)* which showed increased expression of *PML* and *HLA-B* in the cerebral cortices of individuals with ASD (Fig. 6, E and F). This, along with the enrichment of ASD-associated genes among the IFNγ-responding genes detected in this study provides compelling evidence for the relevance of antiviral signalling to ASD pathogenesis.

## Discussion

Epidemiological and animal studies support a role for antiviral inflammatory activation in the aetiology of neurodevelopmental disorders. While the molecular mechanisms that underlie this association are unclear, inflammatory cytokines are thought to play a central role *(11)*. In this study, we used hIPSCs to examine the impact of IFNγ exposure during human neuronal differentiation. Intriguingly, we observed morphological and transcriptomic changes previously associated with neurodevelopmental disorders. We propose a model where IFNγ promotes neurite outgrowth by a PML-dependent long-lasting upregulation of MHCI proteins at neuronal growth cones (Fig. 4J). To our knowledge, the mechanisms through which transient developmental immune activation cause lasting alterations in neuronal phenotypes have not previously been examined in a human system.

IFNγ exposure at the neural progenitor stage induced a robust increase in neurite outgrowth in human neurons. This was measurable as both neurite length and branch number. Our observation is consistent with studies of mouse NPCs *(41, 42)* and a human cancer cell line *(43)*. This study is the first report of this effect in human neurons. Neurite outgrowth is a fundamental stage of neuronal maturation, where neural progenitors extend processes which can later become axons or dendrites. Alterations in neurite outgrowth in the developing brain would be predicted to have implications for neuronal connectivity and ultimately brain function. Increased neurite outgrowth is a consistent finding from studies of hIPSC-neurons derived from individuals with ASD *(25–27)*. Abnormalities have also been observed in hIPSC-neurons from SZ patients *(44)*. Post-mortem studies of individuals diagnosed with ASD and SZ have pointed to abnormal cortical neuron organisation, dendritic arborization and dendritic spine density *(45)*. The macrocephaly present in some ASD cases has been attributed to increased dendrite number and size *(46)*. Interestingly, offspring of pregnant dams exposed to a single gestational dose of poly(I:C) also display altered cortical development, with perturbed dendritic and synaptic development *(6, 11, 21, 24)*. Further evidence comes from large-scale genetic studies of ASD and SZ, which consistently identify genes encoding synaptic proteins and those involved in neuronal maturation *(39, 47)*. Together, these reports highlight the relevance of disturbed neurite outgrowth to the pathophysiology of neurodevelopmental disorders.

A key question we sought to address was how transient immune activation could have a lasting impact on neuronal phenotype. PML nuclear bodies are chromatin associated organelles that play an important role in viral infection response and transcriptional regulation. Moreover, PML has been shown to be involved in neuronal development and function *(22, 23)*. We found that IFNγ established a long-lasting increase in PML body number during the neuronal differentiation of hIPSCs. PML bodies were spatially associated with the site of *HLA-B* transcription and their disruption prevented IFNγ-induced MHCI transcription. Furthermore, PML body disruption blocked IFNγ-induced neurite outgrowth. While this phenomenon has not been examined before in human neurons, interferon-induced transcriptional memory has previously been observed in human cancer cells *(20)*, mouse embryonic fibroblasts and mouse bone-marrow derived macrophages *(19)*. In the latter study, IFNγ pre-treated cells were shown to mount an enhanced antiviral transcriptional response, conferring resistance to viral infection *(19)*. Importantly, PML nuclear bodies were shown to mediate IFNγ-dependent transcriptional memory for the MHC Class II DRA gene in human cancer cells *(20)*. Building on these findings, our data indicate that IFNγ exposure in neural progenitor cells establishes PML-dependent transcriptional memory in MHCI genes that persists through human neuronal differentiation. Remarkably, we also detected a higher number of PML bodies per nucleus in unstimulated hIPSC-NPCs derived from individuals with ASD when compared with controls. This is consistent with the observation that the *PML* gene is upregulated in the post-mortem ASD brain *(15)*.

Genes of the MHCI complex were among the top differentially expressed in both neural progenitors and neurons following IFNγ exposure. They exhibited evidence of transcriptional memory, where expression remained elevated in neurons that received a single IFNγ treatment at the NPC stage. Furthermore, these genes had a heightened response in neurons when primed at the progenitor stage. Using super resolution microscopy, we found that MHCI proteins were enriched in neuronal growth cones. Importantly, disruption of PML body formation prevented both MHCI enrichment in growth cones and IFNγ-induced neurite outgrowth. Moreover, IFNγ did not elicit the same neurite outgrowth phenotype in cells devoid of MHCI at the cell surface. These data indicate that MHCI proteins are involved in IFNγ-induced neurite outgrowth. The role MHCI proteins play in the cellular antiviral response is well established; they present antigens at the cell surface for detection by T cells. In addition, evidence is mounting for non-immune functions in the central nervous system. MHCI proteins have been found in neurons of the developing mammalian brain, where they have been shown to localise to neurites and neuronal growth cones*(16, 31)*. They have been implicated in neurite outgrowth in mouse primary hippocampal neurons *(15)* and synaptic stability in the mouse visual system *(48)*. Intriguingly, gestational poly(I:C) also leads to an enduring increase in MHCI protein in mouse cortical neurites *(21)*. It is noteworthy that the MHC locus has shown the strongest association with SZ in multiple major genome wide association studies *(49, 50)*, however, strong linkage disequilibrium in the region has made identifying the causal variants a challenge. Combined with the existing literature, our results indicate that MHCI proteins are involved in IFNγ-induced neurite outgrowth. Further research is required to determine the mechanisms through which MHCI proteins alter growth cone dynamics.

A growing body of evidence indicates that genetic variants associated with ASD and SZ manifest their risk during critical periods of early brain development *(8, 51–53)*. Alterations in the expression of many risk genes combine to disturb fundamental neurodevelopmental processes involved in neuronal differentiation and maturation *(39, 54)*. In this study, we found that IFNγ exposure during human neuronal differentiation disproportionately altered the expression of genes associated with ASD and SZ. More specifically, risk genes from both disorders were over-represented among genes downregulated by NPCs in response to IFNγ. SZ risk genes were also over-represented among genes upregulated by neurons in response to IFNγ. In total, 66 ASD and 104 SZ risk genes responded to IFNγ in our study with several high-profile candidates among them. One such example is *PTEN*, a tumour suppressor gene considered a high confidence risk gene for ASD *(38)*. The disease associated mutations promote cortical macrocephaly observed in some ASD cases *(55)*. Interestingly, *PTEN* was downregulated in neurons after IFNγ exposure in our study, consistent with the reduced activity caused by ASD-associated variants.

Other outstanding examples include *ZNF804A, FOXP1, TSNARE1* and *TCF4*. It is not clear whether these IFNγ-responding risk genes are involved in the neurite phenotype we describe above. However, similar dysregulation in response to developmental inflammatory stimuli in vivo would be expected to have implications for brain development. The convergence of genetic and environmental risk factors on the same genes is noteworthy and may speak to the association between maternal immune activation and neurodevelopmental disorders.

We also observed a significant overlap between genes that were differentially expressed in response to IFNγ in our model with those found to be dysregulated in post-mortem ASD and SZ brains. The overlapping genes were enriched for genes of the IFNγ signalling pathway and antiviral response genes. The upregulation of immune and inflammatory factors is a consistent finding of transcriptomic studies of ASD and SZ brains *(10)*. While this signal has typically been assumed to be driven by microglia, our results suggest that it may also have a neuronal origin. This observation provides validity to our model as IFNγ alters gene expression in a manner consistent with the dysregulation observed in the brains of patients with these disorders.

In summary, we find that antiviral immune activation in human NPCs induces morphological and transcriptomic changes associated with neurodevelopmental disorders. NPCs are thought to be central to the aetiology of these conditions. Their disruption can have lasting implications for neuronal migration, maturation and function *(56)*. Recently, Schafer and colleagues observed that hIPSC-neurons from individuals with ASD displayed increased neurite outgrowth *(27)*. Importantly, they demonstrated that this phenotype was mediated by aberrant gene expression in NPCs and could be prevented by bypassing the NPC stage, through direct conversion to neurons. In our study, we find that pathological priming of NPCs with IFNγ has a remarkably similar effect. Thus, perturbation of normal gene expression in NPCs, whether genetic or environmental, leads to altered neuronal maturation. The degree to which this phenotype contributes to neurodevelopmental disorders remains an open question. Further investigation is required to determine whether the IFNγ-responding ASD and SZ risk genes contribute to the neurite outgrowth phenotype and whether PML or MHCI mediate perturbed risk gene expression. Nonetheless, our results highlight antiviral signalling as a plausible link between early immune activation and neurodevelopmental disorders. This work provides a framework for future study of immune activation and gene-environment interaction in human neural development.

## Materials and Methods

### Study design

The initial objective was to determine whether IFNγ treatment of hIPSC-NPCs followed by continued differentiation into postmitotic neurons led to recapitulation of morphological characteristics of hIPSC-neurons derived from individuals with ASD. Following confirmation of this, we carried out RNA-sequencing to determine the transcriptional changes brought about by acute treatment, pre-treatment and double-treatment with IFNγ in NPCs and neurons. Further hypotheses were generated to test the requirement for PML nuclear bodies in IFNγ-dependent MHCI transcription and the requirement for PML and MHCI in IFNγ-dependent increased neurite outgrowth. We also carried out gene enrichment analysis to test the hypotheses that IFNγ would disproportionately alter the expression of ASD and SZ risk genes as well as those that are differentially expressed in post-mortem brains of individuals with these disorders. No data were excluded from any dataset. To account for variability between cultures, multiple biological replicates were generated where appropriate and as specified. Cells from a given hIPSC line were considered to be independent biological replicates when they were generated from hIPSC samples with a different passage number. Treatments were carried out on cells within the same biological replicate and paired or grouped statistical analysis was carried out, to limit the impact of this variability on measured outcomes. The numbers of hIPSC lines and biological replicates, sample sizes and statistical tests used are specified in figure legends. Experiment specific parameters are outlined in the methodology section below.

### HiPSC generation and neuralisation and treatment

Participants were recruited and methods carried out in accordance to the ‘Patient iPSCs for Neurodevelopmental Disorders (PiNDs) study’ (REC No 13/LO/1218). Informed consent was obtained from all subjects for participation in the PiNDs study. Ethical approval for the PiNDs study was provided by the NHS Research Ethics Committee at the South London and Maudsley (SLaM) NHS R&D Office. HIPSCs were generated from human hair follicle keratinocytes, derived from three control males with no known psychiatric conditions and three males with diagnosed ASD (Supplementary Table 1). Three control and one ASD hIPSC lines were generated using a polycistronic lentiviral construct co-expressing the four reprogramming transcription factors, OCT4, SOX2, KLF4 and c-MYC, as previously reported *(28)*. Two ASD lines were generated using CytoTune™ Sendai reprogramming kits (Thermo Fischer A16517). HIPSCs reprogramming was validated as previously described *(26, 28, 57, 58)*. Briefly, genome-wide expression profiling using Illumina BeadChip v4 and the bioinformatics tool ‘Pluritest’ was used to confirm hIPSC identity. Pluripotency was established through differentiation into embryoid bodies, followed by immunocytochemistry for markers from the three germ layers. Expression of pluripotency markers, NANOG, OCT4, SSEA4 and TRA1-81 was confirmed by immunocytochemistry. The Illumina Human CytoSNP-12v2.1 BeadChip array and KaryoStudio analysis software (Illumina) were used to assess genome integrity (Supplementary Table 1).

HIPSCs were cultured in StemFlex media (Gibco, A3349401) on 6-well plates (Thermo Fischer: 140675) coated with Geltrex basement membrane matrix (Thermo Fisher, A1413302). Cells were passaged upon reaching 60-70% confluency, by incubation with a calcium chelator, EDTA (Thermo Fisher: 15040-33), followed by detachment with a cell lifter to maintain intact hIPSC colonies.

HIPSCs were differentiated as previously reported *(26)*. HIPSCs were grown until 95-100% confluent then neuralised using a modified dual SMAD inhibition protocol *(59)*. For neuralisation, StemFlex was replaced with a 1:1 mixture of N2 and B27 supplemented medium, made following manufacturer guidelines (Thermo Fisher, 17502048; 17504044) to which SMAD and Wnt inhibitors, (SB431542, 10μM, and Dorsomorphin, 1μM and XAV939, 2μM) were added (hereafter known as N2:B27+++). This was replaced daily until Day (D) 7, when cells were dissociated using Accutase (Life Technologies, A11105-01) and re-plated at a 1:1 ratio into N2:B27 +++, supplemented with Rock inhibitor (Merck, Y27632) to prevent apoptosis. From D8 onwards, SMAD and WNT inhibitors were excluded from medium and cells received daily N2:B27 medium alone. Cells were dissociated and re-plated on D12, D16 and D19 and D21. From D9 a uniform sheet of SOX2 / NESTIN positive cells was observed (Supplementary Fig. 1). From D17 cells were observed to self-organise into neuroepithelial rosettes, expressing apical polarity marker PKCλ with mitotic, PH3-expressing cells located at the apical pole (Fig. S1). On D21 cells were plated into Poly-D-Lysine (5µg/ml Merck, P6407) and laminin (20µg/ml, Merck, L2020) coated plates, in B27 media, supplemented with Notch inhibitor, DAPT, to induce cell cycle exit. From this point onwards βIII-Tubulin expression and neurite extension were observed (Fig S1). Cells received IFNγ (Abcam, AB9659) at a concentration of 25ng/ml on D17-21 and D29-30 and arsenic trioxide, As_2_O_3_ (Trisenox™, Teva) at a concentration of 1µM on D16-21, as specified.

### Embryonic stem cell generation and neuralisation and treatment

The human embryonic stem cell (hESC) line, MasterShef 3 HLA null was cultured as previously described for hIPSCs, but using Laminin LN-521 5µg/ml (BioLamina, Sweden) coated plates. Mastershef 3 HLA null was created using CRISPR Cas9 nickase technology. Two plasmids containing a mutant Cas9D10A nickase plus different reporter genes GFP and dTomato (pSpCas9n(BB)_2A_GFP, pSpcas9n(BB)_2A_dTomato) with different guide RNAs targeting the Beta-2-Microglobulin (B2M) gene were transfected into the parent line, followed by single cell sorting 4 days later for GFP/dTomato double positive cells. Single cells were seeded onto LN-521 coated 96 well plates and colonies picked after 7-10 days. Colonies were screened by PCR for B2M expression, by flow cytometry for MHCI knockout, by T7 mismatch cleavage assay and finally Sanger sequencing of chosen clones. hESCs were expanded and differentiated using identical protocols to hIPSC lines described above.

### *B2M* knock down

*B2M* was knocked down using lentiviral particles containing *B2M*-targeting shRNA (OriGene, TL314543V, Virus A). Non-targeting scramble shRNA from the same kit was used as control. Cells were exposed to the shRNA on D14 for 18 hours at MOI 8. NPCs were treated with IFNγ on D17 for 24h and fixed on D18. These experiments were carried out in biological triplicate for all three control hIPSC lines.

### RNA Sequencing

RNA samples were collected in Trizol Reagent (Thermo Fisher, 15596026) and RNA extraction was performed using an RNEasy kit (Qiagen, 74104). Each experimental condition was repeated in the three control hIPSC lines (Supplementary Table 1). Libraries were prepared using the TruSeq RNA Library Preparation Kit (Illumina, RS-122-2001) by the Wellcome Trust Centre for Human Genetics, University of Oxford. Libraries were pooled and sequenced on an Illumina HiSeq 4000, resulting in an average of 38 million 75 bp paired-end reads per sample. FASTQ files were processed with Trimmomatic *(60)* to quality trim the reads and remove adapters. The reads were then mapped to the human (GRCh38) reference genome using STAR *(61)*. Count matrices were prepared using GenomicAlignments on Bioconductor *(62)* and differential gene expression analysis was carried out with DESeq2 *(63)*. The threshold for statistical significance was Benjamini-Hochberg (BH) adjusted *P* value < 0.05. Genes with an adjusted *P* value of 0 were awarded *padj* = 1 x 10^−308^ for the purpose of the volcano plots. Gene ontology analysis was carried out with DAVID *(64)* for biological process, molecular function and cellular component. Differentially expressed gene lists were tested for enrichment relative to a background of all genes awarded an adjusted *P* value by DESeq2 for that comparison. A BH-adjusted *P* value < 0.05 was the threshold for enrichment. The raw sequencing data can be accessed from www.synapse.org/IFNG.

### Gene enrichment analysis

The ASD risk gene list was from the SFARI Gene database *(38)* and included all 736 genes from category 1-4 of the January 15^th^ 2019 release. The SZ list included the 1,111 risk genes compiled by the PsychENCODE Consortium (PEC) *(39)*. None of the risk genes included in this analysis fall in the MHC region. Differentially expressed genes from post-mortem brains of ASD and SZ patients were taken from the PEC cross-disorder study, which employed 51 ASD, 559 SZ and 936 control post-mortem frontal and temporal cerebral cortex samples *(10)*. Enrichment of the above gene sets in our DEGs was tested using a two tailed Fisher’s exact test in R. BH correction for multiple testing was applied. Up and down-regulated genes were tested separately. Background was controlled for by testing gene frequency within the differentially expressed sets against the frequency in the equivalent tested but non-differentially expressed set (awarded an adjusted *P* value > 0.05 by DESeq2).

### Immunocytochemistry

Cells were washed in phosphate-buffered saline (PBS), fixed in 4% paraformaldyhyde (20 minutes at room temperature), washed three times in PBS and permeabilised and blocked using a solution of 4% Donkey Serum and 0.1% Triton in PBS, for one hour at room temperature (RT). Primary antibodies (Supplementary Table 2) were diluted in blocking solution and applied overnight at 4°C. Following this, cells were washed three times in PBS and appropriate Alexa Fluor secondary antibodies (A-21202, A-21203, A-21206, A-21207, ThermoFisher) were applied in blocking solution for two hours at RT. Cells were then washed three times in PBS and counterstained with Hoechst 33342 (Merck, B2261). F-actin was detected using Alexa Fluor 488-conjugated phalloidin (ActinGreen 488 ReadyProbes, ThermoFischer, R37110), applied in PBS for 30 minutes following secondary antibody incubation and three washes in PBS.

### High Content Cellomic Screening

For the initial neuronal morphology experiments, high content screening was performed using CellInsight™ (Thermo Scientific) operated by HCS Studio Software for automated measurement of neurite outgrowth. Cells were plated at a density of 9000 cells/cm^2^ in PDL and laminin-coated 96 well Nunclon® plates (Merck, P8366), stained for βIII-Tubulin (Tuj1) for detection of both mature and immature neurites and counterstained with Hoechst 33342 to detect nuclei. The analysis pipeline involved initial detection of Hoechst-stained nuclei, detection of cell bodies using nuclei as seeds then detection of neurites originating from cell bodies. The following features were assessed for comprehensive examination of neuronal morphology: neurite total length per cell, neurite average length per cell, neurite count per cell, branch point count per cell. Averages were then taken for each condition within biological replicates. Identical detection parameters were used between conditions to allow direct comparison and paired statistical analysis. Analogous analysis was carried out for the MS3 HLA null hESC experiment with the Opera Phenix high content screening system (Perkin Elmer). Plating and staining were carried out as described above. The Opera Phenix was also used to quantify B2M and MHCI protein levels for the *B2M* shRNAi-mediated knock down and MS3 HLA null hESC knock out studies. In both cases, cells were fixed and stained immediately after IFNγ treatment at D18. Protein expression was measured as average cellular intensity, normalised to no-primary controls.

### Quantitative PCR

RNA samples were collected in Trizol Reagent (Thermo Fisher, 15596026) and RNA extraction was performed using an RNEasy kit (Qiagen, 74104). DNA digest was performed using a TURBO DNA-free kit (Thermo Fischer, AM1907) to remove residual genomic DNA from samples. Reverse transcription to generate cDNA, was performed using SuperScript III (Invitrogen, 18080-044). QPCR was performed using EvaGreen Hot FirePol qPCR Mix Plus (Solis Biodyne, 08-24-00001) in a Biorad Chromo4 PCR detection system. QPCR primer sequences can be found in Supplementary Table 3. Gene expression fold-change between conditions was calculated using the Pfaffl comparative Ct method *(65)*.

### RNA Fluorescence *In Situ* Hybridisation

RNA FISH was carried out using an RNAScope™ double Z probe targeting pre-splicing *HLA-B* RNA (ACD, probe named Hs-HLAB-intron). Hybridisation and signal amplification was carried out using RNAScope™ 2.5 HD Detection Reagent kit (ACD, 322350), following manufacturer instructions. Briefly, cells were grown in 8-well chamber slides (Merck, C7057), fixed in 10% Neutral Buffered Formalin for 30 minutes at RT, washed in PBS and incubated with the probe for 2 hours at 40°C. Slides were washed and signal was amplified through a 6-step amplification process, followed by signal detection with a Fast-red chromogen label.

### Microscopy and image analysis

Confocal images were taken with a Leica SP5 laser scanning confocal microscope, using 405/488/594nm lasers and a 63x (NA 1.4) oil immersion lens (Leica, Wetzlar, Germany), operated through Leica Application Suite Advanced Software (v.2.7.3). Super-resolution images were taken using a Visitech-iSIM module coupled to a Nikon Ti-E microscope with a Nikon 100x 1.49 NA TIRF oil immersion lens (Nikon, Japan), using 405/488/561nm lasers. Super-resolution images were deconvolved to increase contrast and resolution, using a Richardson-Lucy algorithm specific to the iSIM mode of imaging using the supplied NIS-Elements Advanced Research software (Nikon, Japan, v.4.6). Identical laser gain and offset settings were used within each biological replicate to enable direct comparison between conditions.

Semi-automated image analysis was performed using ImageJ software. Gaussian filters were applied to PML and RNAScope images to reduce noise and binary images were generated. Nuclei were used as boundaries within which PML and RNAScope spots were counted. Identical processing parameters were applied across conditions to allow paired or grouped statistical analysis. Numbers of PML bodies per µM^2^ were measured within RNAScope spots, within RNAScope spot perimeter regions and within whole nuclei. RNAScope spot perimeter regions were defined as detected spot perimeters +/- 0.3 µm, generating a ring-shaped region of interest.

DiAna, an Image J Plugin *(36)*, was used to assess the spatial distribution of objects. Binary images were generated from confocal images of PML and RNAScope staining. Centre-to-centre distances were measured from PML spots to RNAScope spots. Binary images of nuclei were used as a bounding box (mask) and randomised shuffle of objects in both channels was carried out 100 times. Centre-to-centre distances in shuffled images were computed and cumulated frequency distributions were calculated for real and simulated distances. This was repeated for 9 images across 3 cell lines. Real and simulated distances within a given image were matched by percentile for paired analysis.

### Data Collection and Statistics

All experiments were carried out using 3 independent IPSC lines derived from 3 unrelated male control or ASD-diagnosed individuals. Where possible multiple biological replicates of each line were used (as specified in figure legends). All data was tested for normal distribution using the D’Agostino and Pearson test before further statistical analysis. Statistical tests used included t tests, Mann-Whitney tests, Wilcoxen matched-pairs test; one-way ANOVA and two-way ANOVAs with Tukey and Sidak’s multiple comparison tests respectively. Statistical analysis was carried out using GraphPad Prism 8.

## Acknowledgements

The study was supported by grants from the Innovative Medicines Initiative Joint Undertaking under grant agreement no. 115300, resources of which are composed of financial contribution from the European Union’s Seventh Framework Programme (FP7/2007-2013) and EFPIA companies’ in kind contribution (JP and DPS). This work was also supported by funds the European Autism Interventions (EU-AIMS), and the Innovative Medicines Initiative Joint Undertaking under grant agreement no. 115300, resources of which are composed of financial contribution from the European Union’s Seventh Framework Programme (FP7/2007-2013) and EFPIA companies’ in kind contribution (JP and DPS). In addition, funds from the Wellcome Trust ISSF Grant (No. 097819) and the King’s Health Partners Research and Development Challenge Fund, a fund administered on behalf of King’s Health Partners by Guy’s and St Thomas’ Charity awarded to DPS; the Brain and Behavior Foundation (formally National Alliance for Research on Schizophrenia and Depression (NARSAD); Grant No. 25957), awarded to DPS, were used to support this study. We thank the Wohl Cellular Imaging Centre (WCIC) at the IoPPN, Kings College, London, for help with microscopy. We further thank Michael Gandal for sharing and permission to use PEC data. Finally, we would like thank Dr. Anthony Vernon (King’s College London), Dr. Nichloas Bray (Cardiff University) and Prof. Rosa Sorrentino (Sapienza University of Rome) for critical discussions on this manuscript.

## Notes

### Competing Interest Statement

The authors have declared no competing interest.

## References

1. M. L. Estes, A. K. Mcallister, Maternal immune activation: Implications for neuropsychiatric disorders, 353 (2016), doi: 10.1126/science.aag3194.

2. A. S. Brown, M. D. Begg, S. Gravenstein, C. A. Schaefer, R. J. Wyatt, Serologic Evidence of Prenatal Influenza in the Etiology of Schizophrenia, Arch Gen Psychiatry 61, 774–780 (2004).

3. M. Byrne, E. Agerbo, B. Bennedsen, W. W. Eaton, P. B. Mortensen, Obstetric conditions and risk of first admission with schizophrenia: A Danish national register based study, Schizophr. Res. 97, 51–59 (2007).

4. H. Ó. Atladóttir, P. Thorsen, L. Østergaard, D. E. Schendel, S. Lemcke, M. Abdallah, E. T. Parner, Maternal infection requiring hospitalization during pregnancy and autism spectrum disorders, J. Autism Dev. Disord. 40, 1423–1430 (2010).

5. H. Ó. Atladóttir, T. B. Henriksen, D. E. Schendel, E. T. Parner, Autism After Infection, Febrole Episodes, and Antibiotic Use During Pregancy: An Exploration Study, Pediatrics 130, 1447–1454 (2012).

6. U. Meyer, Prenatal Poly(I:C) exposure and other developmental immune activation models in rodent systems, Biol. Psychiatry 75, 307–315 (2014).

7. I. Voineagu, X. Wang, P. Johnston, J. K. Lowe, Y. Tian, S. Horvath, J. Mill, R. M. Cantor, B. J. Blencowe, D. H. Geschwind, Transcriptomic analysis of autistic brain reveals convergent molecular pathology., Nature 474, 380–4 (2011).

8. N. N. Parikshak, R. Luo, A. Zhang, H. Won, J. K. Lowe, V. Chandran, S. Horvath, D. H. Geschwind, Integrative functional genomic analyses implicate specific molecular pathways and circuits in autism, Cell 155, 1008–1021 (2013).

9. S. Gupta, S. E. Ellis, F. N. Ashar, A. Moes, J. S. Bader, J. Zhan, A. B. West, D. E. Arking, Transcriptome analysis reveals dysregulation of innate immune response genes and neuronal activity-dependent genes in autism., Nat. Commun. 5, 5748 (2014).

10. M. J. Gandal, P. Zhang, E. Hadjimichael, R. L. Walker, C. Chen, S. Liu, H. Won, H. Van Bakel, M. Varghese, Y. Wang, A. W. Shieh, J. Haney, S. Parhami, J. Belmont, M. Kim, P. M. Losada, Z. Khan, J. Mleczko, Y. Xia, R. Dai, D. Wang, Y. T. Yang, M. Xu, K. Fish, P. R. Hof, J. Warrell, D. Fitzgerald, K. White, A. E. Jaffe, M. A. Peters, M. Gerstein, C. Liu, L. M. Iakoucheva, D. Pinto, D. H. Geschwind, Transcriptome-wide isoform-level dysregulation in ASD, schizophrenia, and bipolar disorder, Science 362 (2018), doi: 10.1126/science.aat8127.

11. P. A. Garay, E. Y. Hsiao, P. H. Patterson, A. K. McAllister, Maternal immune activation causes age- and region-specific changes in brain cytokines in offspring throughout development, Brain. Behav. Immun. 31, 54–68 (2013).

12. W. M. Schneider, M. D. Chevillotte, C. M. Rice, Interferon-stimulated genes: a complex web of host defenses., Annu. Rev. Immunol. 32, 513–45 (2014).

13. P. E. Goines, L. A. Croen, D. Braunschweig, C. K. Yoshida, J. Grether, R. Hansen, M. Kharrazi, P. Ashwood, J. Van de Water, Increased midgestational IFN-γ, IL-4 and IL-5 in women bearing a child with autism: A case-control study., Mol. Autism 2, 13 (2011).

14. L. S. Heuer, L. A. Croen, K. L. Jones, C. K. Yoshida, R. L. Hansen, R. Yolken, O. Zerbo, G. DeLorenze, M. Kharrazi, P. Ashwood, J. Van de Water, An Exploratory Examination of Neonatal Cytokines and Chemokines as Predictors of Autism Risk: The Early Markers for Autism Study, Biol. Psychiatry, 1–10 (2019).

15. T. Bilousova, H. Dang, W. Xu, S. Gustafson, Y. Jin, L. Wickramasinghe, T. Won, G. Bobarnac, B. Middleton, J. Tian, D. L. Kaufman, Major histocompatibility complex class I molecules modulate embryonic neuritogenesis and neuronal polarization, J. Neuroimmunol. 247, 1–8 (2012).

16. L. A. Needleman, X.-B. Liu, F. El-Sabeawy, E. G. Jones, A. K. McAllister, MHC class I molecules are present both pre- and postsynaptically in the visual cortex during postnatal development and in adulthood, Proc. Natl. Acad. Sci. 107, 16999–17004 (2010).

17. C. S. Nicolas, M. Amici, Z. A. Bortolotto, A. Doherty, Z. Csaba, A. Fafouri, P. Dournaud, P. Gressens, G. L. Collingridge, S. Peineau, The role of JAK-STAT signaling within the CNS, Jak-Stat 2, e22925 (2013).

18. A. J. Filiano, Y. Xu, N. J. Tustison, R. L. Marsh, W. Baker, I. Smirnov, S. D. Turner, Z. Weng, S. N. Peerzade, C. C. Overall, P. Sachin, neuronal connectivity and social behaviour, Nature 535, 425–429 (2016).

19. R. Kamada, W. Yang, Y. Zhang, M. C. Patel, Y. Yang, R. Ouda, A. Dey, Interferon stimulation creates chromatin marks and establishes transcriptional memory, Proc. Natl. Acad. Sci. U. S. A. (2018), doi: 10.1073/pnas.1720930115.

20. M. Gialitakis, P. Arampatzi, T. Makatounakis, J. Papamatheakis, Gamma interferon-dependent transcriptional memory via relocalization of a gene locus to PML nuclear bodies., Mol. Cell. Biol. 30, 2046–56 (2010).

21. B. M. Elmer, M. L. Estes, S. L. Barrow, A. K. McAllister, MHCI requires MEF2 transcription factors to negatively regulate synapse density during development and in disease., J. Neurosci. 33, 13791–804 (2013).

22. E. Korb, C. L. Wilkinson, R. N. Delgado, K. L. Lovero, S. Finkbeiner, Arc in the nucleus regulates PML-dependent GluA1 transcription and homeostatic plasticity., Nat. Neurosci. 16, 874–83 (2013).

23. T. Regad, C. Bellodi, P. Nicotera, P. Salomoni, The tumor suppressor Pml regulates cell fate in the developing neocortex., Nat. Neurosci. 12, 132–140 (2009).

24. Y. Li, G. Missig, B. C. Finger, S. M. Landino, A. J. Alexander, E. L. Mokler, J. O. Robbins, Y. Manasian, W. Kim, K. S. Kim, C. J. McDougle, W. A. Carlezon, V. Y. Bolshakov, Maternal and early postnatal immune activation produce dissociable effects on neurotransmission in mPFC–amygdala circuits, J. Neurosci. 38, 3358–3372 (2018).

25. A. Deshpande, S. Yadav, D. Q. Dao, Z. Y. Wu, K. C. Hokanson, M. K. Cahill, A. P. Wiita, Y. N. Jan, E. M. Ullian, L. A. Weiss, Cellular Phenotypes in Human iPSC-Derived Neurons from a Genetic Model of Autism Spectrum Disorder, Cell Rep. 21, 2678–2687 (2017).

26. A. Kathuria, P. Nowosiad, R. Jagasia, S. Aigner, R. D. Taylor, L. C. Andreae, N. J. F. Gatford, W. Lucchesi, D. P. Srivastava, J. Price, Stem cell-derived neurons from autistic individuals with SHANK3 mutation show morphogenetic abnormalities during early development, Mol. Psychiatry 23, 735–746 (2018).

27. S. T. Schafer, A. C. M. Paquola, S. Stern, D. Gosselin, M. Ku, M. Pena, T. J. M. Kuret, M. Liyanage, A. A. Mansour, B. N. Jaeger, M. C. Marchetto, C. K. Glass, J. Mertens, F. H. Gage, Pathological priming causes developmental gene network heterochronicity in autistic subject-derived neurons, Nat. Neurosci. 22 (2019), doi: 10.1038/s41593-018-0295-x.

28. G. Cocks, S. Curran, P. Gami, D. Uwanogho, A. R. Jeffries, A. Kathuria, W. Lucchesi, V. Wood, R. Dixon, C. Ogilvie, T. Steckler, J. Price, The utility of patient specific induced pluripotent stem cells for the modelling of Autistic Spectrum Disorders, Psychopharmacology (Berl). 231, 1079–1088 (2014).

29. S. M. Saini, S. G. Mancuso, M. S. Mostaid, C. Liu, C. Pantelis, I. P. Everall, C. A. Bousman, Meta-analysis supports GWAS-implicated link between GRM3 and schizophrenia risk, Transl. Psychiatry 7, e1196 (2017).

30. D. W. Volk, T. Matsubara, S. Li, E. J. Sengupta, D. Georgiev, Y. Minabe, A. Sampson, T. Hashimoto, D. A. Lewis, Deficits in transcriptional regulators of cortical parvalbumin neurons in schizophrenia, Am. J. Psychiatry 169, 1082–1091 (2012).

31. M. W. Glynn, B. M. Elmer, P. A. Garay, X. Liu, A. Leigh, F. El-sabeawy, A. K. Mcallister, Establishment of Cortical Connections, 14, 442–451 (2012).

32. N. Masroori, N. Merindol, L. Berthoux, The interferon-induced antiviral protein PML (TRIM19) promotes the restriction and transcriptional silencing of lentiviruses in a context-specific, isoform-specific fashion, Retrovirology 13, 1–17 (2016).

33. X. Zhang, X. Yan, Z. Zhou, F. Yang, Z. Wu, H. Sun, W. Liang, A. Song, V. Lamelland-Breitenbach, M. Jeanne, Q. Zhang, H. Yang, Q. Huang, G. Zhou, J. Tong, Y. Zhang, J. Wu, H. Hu, H. de Thé, S. Chen, Z. Chen, Arsenic trioxide controls the fate of the PML-RARα Oncoprotein by directly binding PML, Science 328, 240–243 (2010).

34. T. Alpert, L. Herzel, K. M. Neugebauer, Perfect timing: splicing and transcription rates in living cells, Wiley Interdiscip. Rev. RNA 8, 1–18 (2017).

35. Y. Brody, Y. Shav-Tal, Transcription and splicing: When the Twain meet, Transcription 2, 222–226 (2011).

36. J. F. Gilles, M. Dos Santos, T. Boudier, S. Bolte, N. Heck, DiAna, an ImageJ tool for object-based 3D co-localization and distance analysis, Methods 115, 55–64 (2017).

37. D. Wang, Y, Quan, Q. Yan, J. E. Morales, R. A. Wetsel, Targeted Disruption of the β2-Microglobulin Gene Minimizes the Immunogenicity of Human Embryonic Stem Cells, Stem Cells Translational Medicine, 1234–1245 (2015).

38. B. S. Abrahams, D. E. Arking, D. B. Campbell, H. C. Mefford, E. M. Morrow, L. A. Weiss, I. Menashe, T. Wadkins, S. Banerjee-basu, A. Packer, SFARI Gene 2.0: knowledgebase for the autism spectrum disorders (ASDs), Mol. Autism, 36 (2013).

39. D. Wang, S. Liu, J. Warrell, H. Won, X. Shi, F. C. P. Navarro, D. Clarke, M. Gu, P. Emani, Y. T. Yang, X. Min, M. J. Gandal, S. Lou, J. Zhang, J. J. Park, C. Yan, S. KyongRhie, K. Manakongtreecheep, H. Zhou, A. Aparna Natha, M. Peters, E. Mattei, D. Fitzgerald, T. Brunetti, J. Moore, Y. Jiang, K. Girdhar, G. E. Hoffman, S. Kalayci, Z. H. Gümüs, G. E. Crawford, P. Roussos, S. Akbarian, A. E. Jaffe, K. P. White, Z. Weng, N. Sestan, D. H. Geschwind, J. A. Knowles, M. B. Gerstein, Comprehensive functional genomic resource and integrative model for the human brain, Science 362 (2018), doi: 10.1126/science.aat8464.

40. M. V. Lombardo, H. M. Moon, J. Su, T. D. Palmer, E. Courchesne, T. Pramparo, Maternal immune activation dysregulation of the fetal brain transcriptome and relevance to the pathophysiology of autism spectrum disorder, Mol. Psychiatry 23, 1001–1013 (2018).

41. G. Wong, Y. Goldshmit, A. M. Turnley, Interferon-γ but not TNFα promotes neuronal differentiation and neurite outgrowth of murine adult neural stem cells, Exp. Neurol. 187, 171–177 (2004).

42. S. J. Kim, T. G. Son, K. Kim, H. R. Park, M. P. Mattson, J. Lee, Interferon-γ promotes differentiation of neural progenitor cells via the JNK pathway, Neurochem. Res. 32, 1399–1406 (2007).

43. J. H. Song, X. W. Chen, D. K. Song, P. Wang, A. Shuaib, C. Hao, Interferon γ induces neurite outgrowth by up-regulation of p35 neuron-specific cyclin-dependent kinase 5 activator via activation of ERK1/2 pathway, J. Biol. Chem. 280, 12896–12901 (2005).

44. K. J. Brennand, A. Simone, J. Jou, C. Gelboin-Burkhart, N. Tran, S. Sangar, Y. Li, Y. Mu, G. Chen, D. Yu, S. McCarthy, J. Sebat, F. H. Gage, Modelling schizophrenia using human induced pluripotent stem cells., Nature 473, 221–5 (2011).

45. V. A. Kulkarni, B. L. Firestein, The dendritic tree and brain disorders, Mol. Cell. Neurosci. 50, 10–20 (2012).

46. Y. N. Jan, L. Y. Jan, Branching out: Mechanisms of dendritic arborization, Nat. Rev. Neurosci. 11, 316–328 (2010).

47. J. Gilbert, H. Y. Man, Fundamental elements in autism: From neurogenesis and neurite growth to synaptic plasticity, Front. Cell. Neurosci. 11, 1–25 (2017).

48. J. Syken, T. GrandPre, P. O. Kanold, C. J. Shatz, PirB Restricts Ocular-Dominance Plasticity in Visual Cortex, Science (80-.). 313, 1795 LP – 1800 (2006).

49. S. Ripke, B. M. Neale, A. Corvin, J. T. R. Walters, K.-H. Farh, P. a. Holmans, P. Lee, B. Bulik-Sullivan, D. a. Collier, H. Huang, T. H. Pers, I. Agartz, E. Agerbo, M. Albus, M. Alexander, F. Amin, S. a. Bacanu, M. Begemann, R. a. Belliveau Jr, J. Bene, S. E. Bergen, E. Bevilacqua, T. B. Bigdeli, D. W. Black, R. Bruggeman, N. G. Buccola, R. L. Buckner, W. Byerley, W. Cahn, G. Cai, D. Campion, R. M. Cantor, V. J. Carr, N. Carrera, S. V. Catts, K. D. Chambert, R. C. K. Chan, R. Y. L. Chen, E. Y. H. Chen, W. Cheng, E. F. C. Cheung, S. Ann Chong, C. Robert Cloninger, D. Cohen, N. Cohen, P. Cormican, N. Craddock, J. J. Crowley, D. Curtis, M. Davidson, K. L. Davis, F. Degenhardt, J. Del Favero, D. Demontis, D. Dikeos, T. Dinan, S. Djurovic, G. Donohoe, E. Drapeau, J. Duan, F. Dudbridge, N. Durmishi, P. Eichhammer, J. Eriksson, V. Escott-Price, L. Essioux, A. H. Fanous, M. S. Farrell, J. Frank, L. Franke, R. Freedman, N. B. Freimer, M. Friedl, J. I. Friedman, M. Fromer, G. Genovese, L. Georgieva, I. Giegling, P. Giusti-Rodríguez, S. Godard, J. I. Goldstein, V. Golimbet, S. Gopal, J. Gratten, L. de Haan, C. Hammer, M. L. Hamshere, M. Hansen, T. Hansen, V. Haroutunian, A. M. Hartmann, F. a. Henskens, S. Herms, J. N. Hirschhorn, P. Hoffmann, A. Hofman, M. V. Hollegaard, D. M. Hougaard, M. Ikeda, I. Joa, A. Julià, R. S. Kahn, L. Kalaydjieva, S. Karachanak-Yankova, J. Karjalainen, D. Kavanagh, M. C. Keller, J. L. Kennedy, A. Khrunin, Y. Kim, J. Klovins, J. a. Knowles, B. Konte, V. Kucinskas, Z. Ausrele Kucinskiene, H. Kuzelova-Ptackova, A. K. Kähler, C. Laurent, J. Lee Chee Keong, S. Hong Lee, S. E. Legge, B. Lerer, M. Li, T. Li, K.-Y. Liang, J. Lieberman, S. Limborska, C. M. Loughland, J. Lubinski, J. Lönnqvist, M. Macek Jr, P. K. E. Magnusson, B. S. Maher, W. Maier, J. Mallet, S. Marsal, M. Mattheisen, M. Mattingsdal, R. W. McCarley, C. McDonald, A. M. McIntosh, S. Meier, C. J. Meijer, B. Melegh, I. Melle, R. I. Mesholam-Gately, A. Metspalu, P. T. Michie, L. Milani, V. Milanova, Y. Mokrab, D. W. Morris, O. Mors, K. C. Murphy, R. M. Murray, I. Myin-Germeys, B. Müller-Myhsok, M. Nelis, I. Nenadic, D. a. Nertney, G. Nestadt, K. K. Nicodemus, L. Nikitina-Zake, L. Nisenbaum, A. Nordin, E. O’Callaghan, C. O’Dushlaine, F. A. O’Neill, S.-Y. Oh, A. Olincy, L. Olsen, J. Van Os, P. Endophenotypes International Consortium, C. Pantelis, G. N. Papadimitriou, S. Papiol, E. Parkhomenko, M. T. Pato, T. Paunio, M. Pejovic-Milovancevic, D. O. Perkins, O. Pietiläinen, J. Pimm, A. J. Pocklington, J. Powell, A. Price, A. E. Pulver, S. M. Purcell, D. Quested, H. B. Rasmussen, A. Reichenberg, M. a. Reimers, A. L. Richards, J. L. Roffman, P. Roussos, D. M. Ruderfer, V. Salomaa, A. R. Sanders, U. Schall, C. R. Schubert, T. G. Schulze, S. G. Schwab, E. M. Scolnick, R. J. Scott, L. J. Seidman, J. Shi, E. Sigurdsson, T. Silagadze, J. M. Silverman, K. Sim, P. Slominsky, J. W. Smoller, H.-C. So, C. a. Spencer, E. a. Stahl, H. Stefansson, S. Steinberg, E. Stogmann, R. E. Straub, E. Strengman, J. Strohmaier, T. Scott Stroup, M. Subramaniam, J. Suvisaari, D. M. Svrakic, J. P. Szatkiewicz, E. Söderman, S. Thirumalai, D. Toncheva, S. Tosato, J. Veijola, J. Waddington, D. Walsh, D. Wang, Q. Wang, B. T. Webb, M. Weiser, D. B. Wildenauer, N. M. Williams, S. Williams, S. H. Witt, A. R. Wolen, E. H. M. Wong, B. K. Wormley, H. Simon Xi, C. C. Zai, X. Zheng, F. Zimprich, N. R. Wray, K. Stefansson, P. M. Visscher, W. Trust Case-Control Consortium, R. Adolfsson, O. a. Andreassen, D. H. R. Blackwood, E. Bramon, J. D. Buxbaum, A. D. Børglum, S. Cichon, A. Darvasi, E. Domenici, H. Ehrenreich, T. Esko, P. V. Gejman, M. Gill, H. Gurling, C. M. Hultman, N. Iwata, A. V. Jablensky, E. G. Jönsson, K. S. Kendler, G. Kirov, J. Knight, T. Lencz, D. F. Levinson, Q. S. Li, J. Liu, A. K. Malhotra, S. a. McCarroll, A. McQuillin, J. L. Moran, P. B. Mortensen, B. J. Mowry, M. M. Nöthen, R. a. Ophoff, M. J. Owen, A. Palotie, C. N. Pato, T. L. Petryshen, D. Posthuma, M. Rietschel, B. P. Riley, D. Rujescu, P. C. Sham, P. Sklar, D. St Clair, D. R. Weinberger, J. R. Wendland, T. Werge, M. J. Daly, P. F. Sullivan, M. C. O’Donovan, Biological insights from 108 schizophrenia-associated genetic loci, Nature (2014), doi: 10.1038/nature13595.

50. J. Shi, D. F. Levinson, J. Duan, A. R. Sanders, Y. Zheng, I. Péer, F. Dudbridge, P. A. Holmans, A. S. Whittemore, B. J. Mowry, A. Olincy, F. Amin, C. R. Cloninger, J. M. Silverman, N. G. Buccola, W. F. Byerley, D. W. Black, R. R. Crowe, J. R. Oksenberg, D. B. Mirel, K. S. Kendler, R. Freedman, P. V. Gejman, Common variants on chromosome 6p22.1 are associated with schizophrenia, Nature 460, 753–757 (2009).

51. A. J. Willsey, S. J. Sanders, M. Li, S. Dong, A. T. Tebbenkamp, R. A. Muhle, S. K. Reilly, L. Lin, S. Fertuzinhos, J. A. Miller, M. T. Murtha, C. Bichsel, W. Niu, J. Cotney, A. G. Ercan-Sencicek, J. Gockley, A. R. Gupta, W. Han, X. He, E. J. Hoffman, L. Klei, J. Lei, W. Liu, L. Liu, C. Lu, X. Xu, Y. Zhu, S. M. Mane, E. S. Lein, Wei, J. P. Noonan, K. Roeder, B. Devlin, N. Sestan, M. W. State, XCoexpression networks implicate human midfetal deep cortical projection neurons in the pathogenesis of autism, Cell 155, 997–1007 (2013).

52. M. J. Hill, J. G. Donocik, R. a Nuamah, C. a Mein, R. Sainz-Fuertes, N. J. Bray, Transcriptional consequences of schizophrenia candidate miR-137 manipulation in human neural progenitor cells., Schizophr. Res. 153, 225–30 (2014).

53. A. E. Jaffe, R. E. Straub, J. H. Shin, R. Tao, Y. Gao, L. Collado-Torres, T. Kam-Thong, H. S. Xi, J. Quan, Q. Chen, C. Colantuoni, W. S. Ulrich, B. J. Maher, A. Deep-Soboslay, A. J. Cross, N. J. Brandon, J. T. Leek, T. M. Hyde, J. E. Kleinman, D. R. Weinberger, Developmental and genetic regulation of the human cortex transcriptome illuminate schizophrenia pathogenesis, Nat. Neurosci. 21, 1117–1125 (2018).

54. J. Grove, S. Ripke, T. D. Als, M. Mattheisen, R. K. Walters, H. Won, J. Pallesen, E. Agerbo, O. A. Andreassen, R. Anney, S. Awashti, R. Belliveau, F. Bettella, J. D. Buxbaum, J. Bybjerg-Grauholm, M. Bækvad-Hansen, F. Cerrato, K. Chambert, J. H. Christensen, C. Churchhouse, K. Dellenvall, D. Demontis, S. De Rubeis, B. Devlin, S. Djurovic, A. L. Dumont, J. I. Goldstein, C. S. Hansen, M. E. Hauberg, M. V Hollegaard, S. Hope, D. P. Howrigan, H. Huang, C. M. Hultman, L. Klei, J. Maller, J. Martin, A. R. Martin, J. L. Moran, M. Nyegaard, T. Nærland, D. S. Palmer, A. Palotie, C. B. Pedersen, M. G. Pedersen, T. dPoterba, J. B. Poulsen, B. S. Pourcain, P. Qvist, K. Rehnström, A. Reichenberg, J. Reichert, E. B. Robinson, K. Roeder, P. Roussos, E. Saemundsen, S. Sandin, F. K. Satterstrom, G. Davey Smith, H. Stefansson, S. Steinberg, C. R. Stevens, P. F. Sullivan, P. Turley, G. B. Walters, X. Xu, A. S. D. W. G. of the P. G. Consortium, BUPGEN, M. D. D. W. G. of the P. G. Consortium, 23andMe Research Team, K. Stefansson, D. H. Geschwind, M. Nordentoft, D. M. Hougaard, T. Werge, O. Mors, P. B. Mortensen, B. M. Neale, M. J. Daly, A. D. Børglum, Identification of common genetic risk variants for autism spectrum disorder, Nat. Genet. 51, 431–444 (2019).

55. A. Orrico, L. Galli, S. Buoni, A. Orsi, G. Vonella, V. Sorrentino, Novel PTEN mutations in neurodevelopmental disorders and macrocephaly, Clin. Genet. 75, 195–198 (2009).

56. R. Stoner, M. L. Chow, M. P. Boyle, S. M. Sunkin, P. R. Mouton, S. Roy, A. Wynshaw-Boris, S. a Colamarino, E. S. Lein, E. Courchesne, Patches of disorganization in the neocortex of children with autism., N. Engl. J. Med. 370, 1209–19 (2014).

57. C. Shum, L. Dutan, E. Annuario, K. Warre-Cornish, S. Taylor, R. Taylor, L. Andreae, N. J. Buckley, J. Price, S. Bhattacharyya, D. P. Srivastava, Δ9-tetrahydrocannabinol negatively regulates neurite outgrowth and Akt signaling in hiPSC-derived cortical neurons., bioRxiv (2018), doi: 10.1101/440909.

58. D. Adhya, V. Swarup, R. Nagy, C. Shum, P. Nowosiad, K. M. Jozwik, I. Lee, D. Skuse, F. A. Flinter, G. McAlonan, M. A. Mendez, J. Horder, D. Murphy, D. H. Geschwind, J. Price, J. Carroll, D. P. Srivastava, S. Baron-Cohen, Atypical neurogenesis and excitatory-inhibitory progenitor generation in induced pluripotent stem cell (iPSC) from autistic individuals, bioRxiv (2018). doi: https://doi.org/10.1101/349415

59. Y. Shi, P. Kirwan, F. J. Livesey, Directed differentiation of human pluripotent stem cells to cerebral cortex neurons and neural networks., Nat. Protoc. 7, 1836–46 (2012).

60. A. M. Bolger, M. Lohse, B. Usadel, Trimmomatic: A flexible trimmer for Illumina sequence data, Bioinformatics 30, 2114–2120 (2014).

61. A. Dobin, C. A. Davis, F. Schlesinger, J. Drenkow, C. Zaleski, S. Jha, P. Batut, M. Chaisson, T. R. Gingeras, STAR: Ultrafast universal RNA-seq aligner, Bioinformatics 29, 15–21 (2013).

62. M. Lawrence, W. Huber, H. Pagès, P. Aboyoun, M. Carlson, R. Gentleman, M. T. Morgan, V. J. Carey, Software for Computing and Annotating Genomic Ranges, PLoS Comput. Biol. 9, 1–10 (2013).

63. M. I. Love, W. Huber, S. Anders, Moderated estimation of fold change and dispersion for RNA-seq data with DESeq2., Genome Biol. 15, 550 (2014).

64. D. W. Huang, B. T. Sherman, R. A. Lempicki, Systematic and integrative analysis of large gene lists using DAVID bioinformatics resources., Nat. Protoc. 4, 44–57 (2009).

65. M. W. Pfaffl, A new mathematical model for relative quantification in real-time RT-PCR, Nucleic Acids Res. 29, 45e – 45 (2001).

