## Supplemental Material for "Antiviral signalling in human IPSC-derived neurons recapitulates neurodevelopmental disorder phenotypes" (from another version of this preprint: version 2, 2020-04-21)


Katherine Warre-Cornish[1,2]†, Leo Perfect[1,2]†, Roland Nagy[1,2], Rodrigo R.R. Duarte[3,4], Matthew J. Reid[1,2], Pooja Raval[1,2], Annett Mueller[5], Amanda L. Evans[5], Amalie Couch[1,2], Cédric Ghevaert[5], Grainne McAlonan[2,6], Eva Loth[2,6], Declan Murphy[2,6], Timothy R. Powell[3,4], Anthony C. Vernon[1,2], Deepak P. Srivastava[1,2]*, Jack Price[1,2,7]*

[1]Department of Basic and Clinical Neuroscience, The Maurice Wohl Clinical Neuroscience Institute, Institute of Psychiatry Psychology and Neuroscience, King's College London, London, SE5 8AF, UK; [2]MRC Centre for Neurodevelopmental Disorders, King's College London, London, UK; [3]Social, Genetic & Developmental Psychiatry Centre, Institute of Psychiatry, Psychology & Neuroscience, King's College London, London, UK; [4]Division of Infectious Diseases, Weill Cornell Medicine, Cornell University, New York, NY, USA; [5]Wellcome-MRC Cambridge Stem Cell Institute; [6]Department of Forensic and Neurodevelopmental Sciences, King's College London, London, UK; [7]National Institute for Biological Standards and Control, South Mimms, UK.

† = These authors contributed equally to this work


### SUPPLEMENTARY MATERIAL

### Supplementary Figures

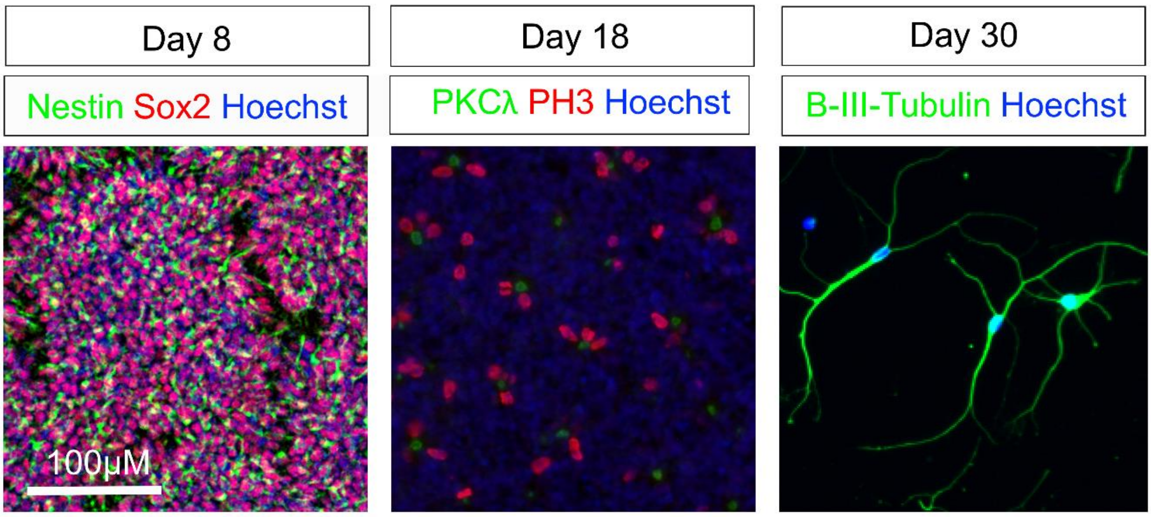


**Supplementary Figure 1: Characterisation of hiPSC neuronal differentiation.** Representative confocal images of neural progenitor cells (NPCs), neural rosettes (also composed of NPCs) and neuronal markers at D8, D18 and D30 during hiPSC neuronal differentiation.

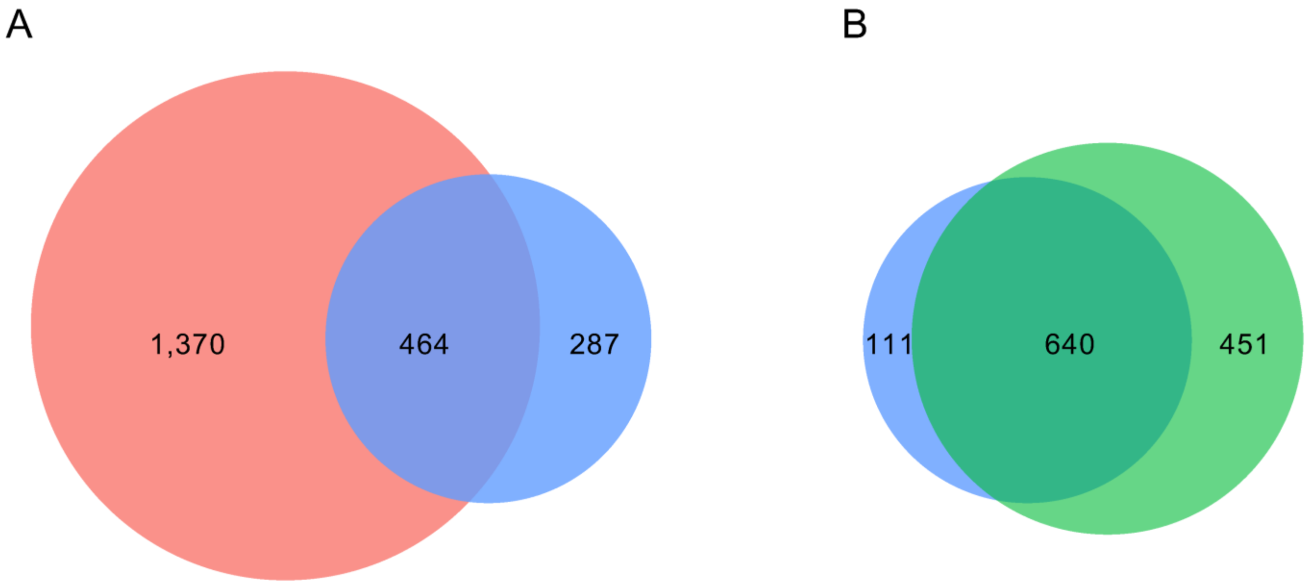


**Supplementary Figure 2: Area proportional Venn diagrams of DEG overlap.** Visualisation of overlapping DEGs from **(A)** 18U vs 18T (red) and 30UU vs 30UT (blue) and **(B)** 30UU vs 30UT (blue) and 30UU vs 30TT (green) conditions.

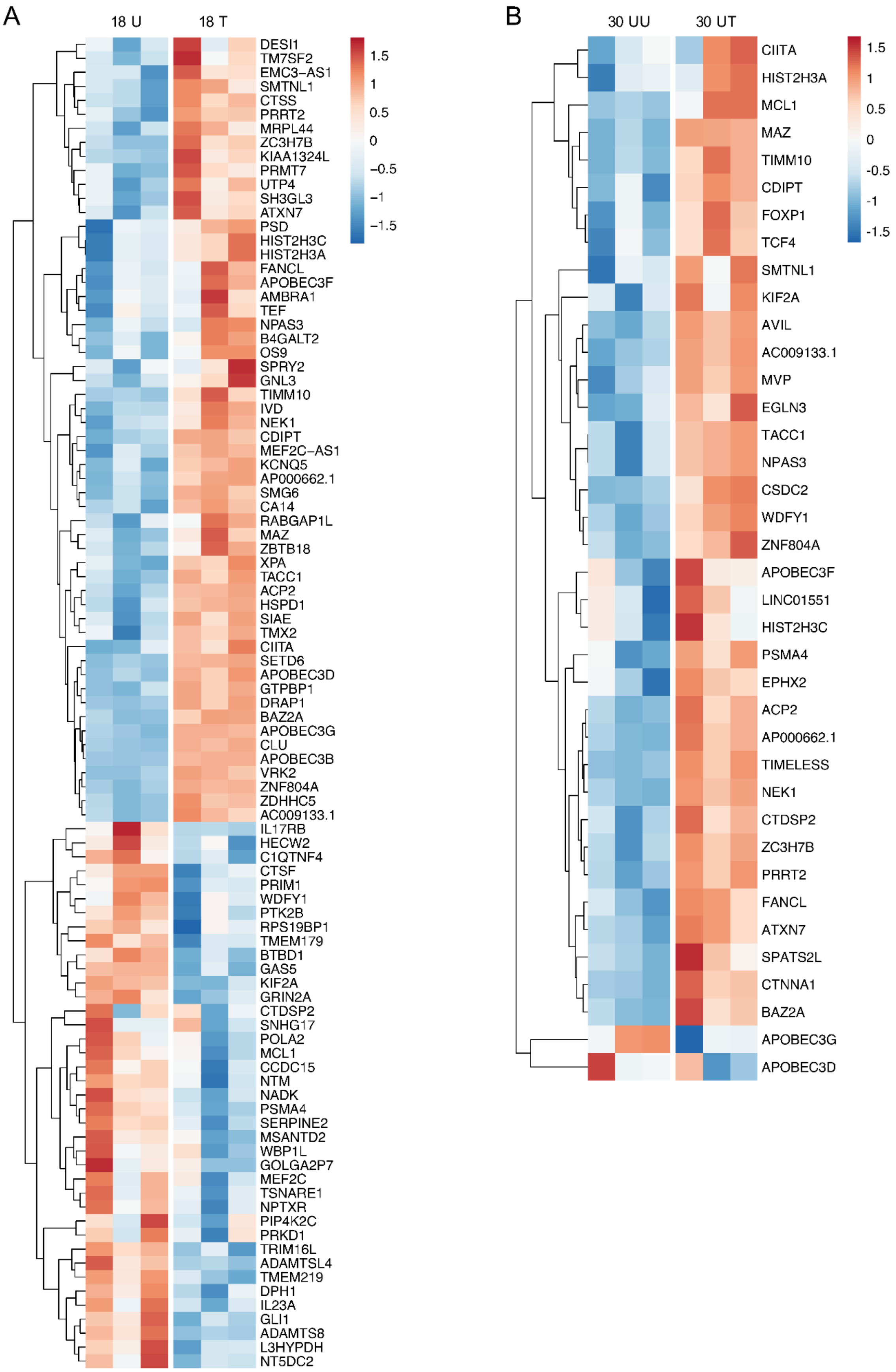


**Supplementary Figure 3: Relative expression of IFNγ-responding SZ risk genes.** Heatmaps illustrating the SZ risk genes from the PsychENCODE gene set that were differentially expressed in response to IFNγ in NPCs (A) and neurons (B). Expression is scaled by row.

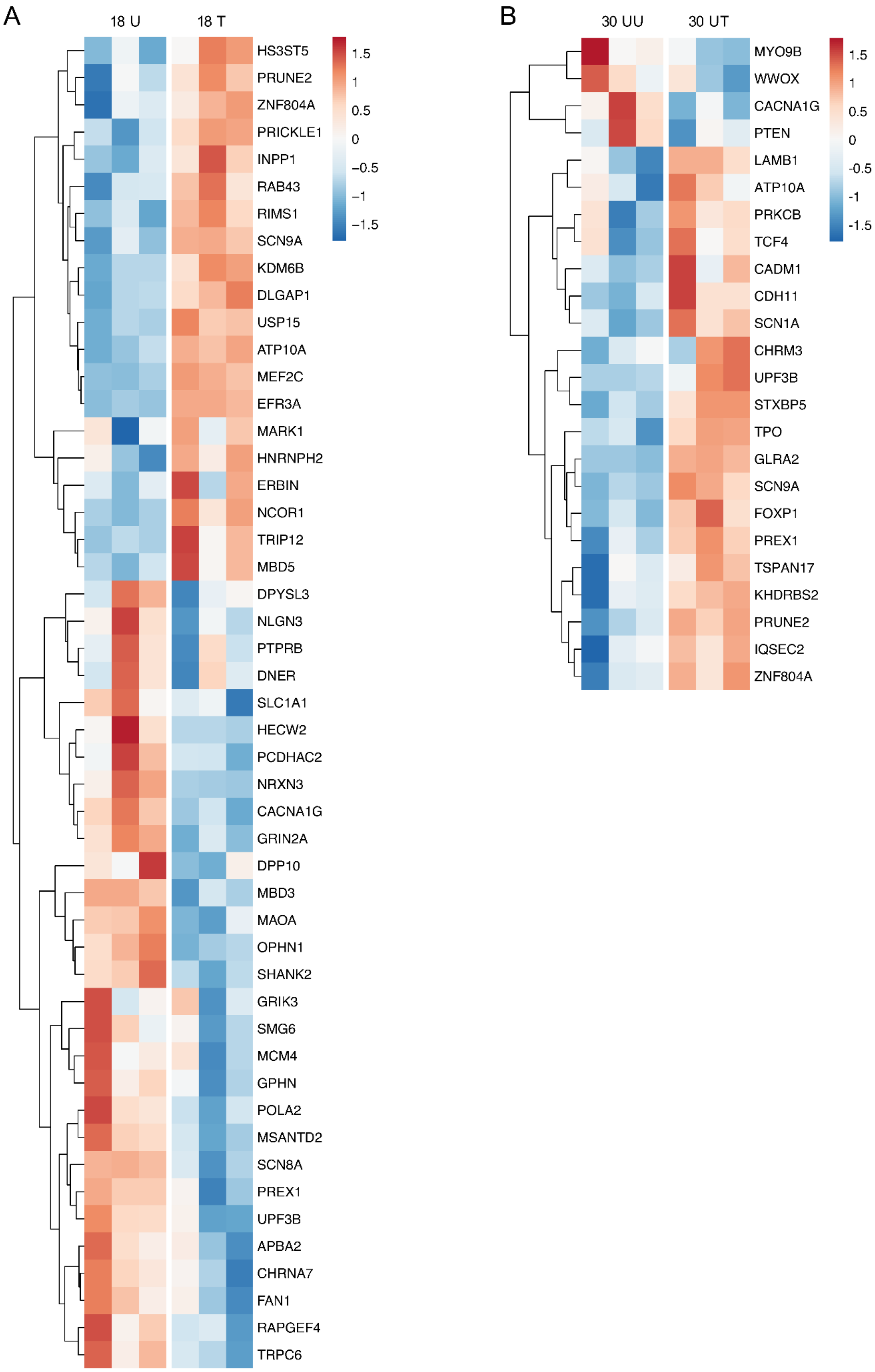


**Supplementary Figure 4: Relative expression of IFNγ-responding ASD risk genes.** Heatmaps illustrating the ASD risk genes from the SFARI database, category 1-4, that were differentially expressed in response to IFNγ in NPCs (A) and neurons (B). Expression is scaled by row.

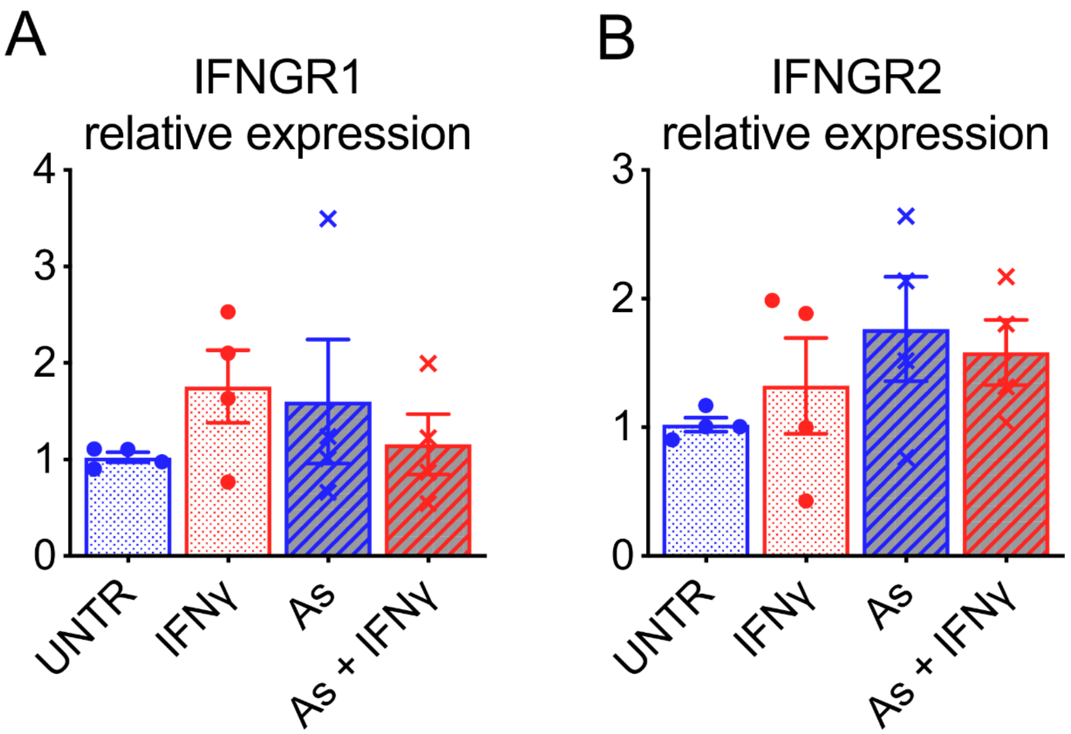


**Supplementary Figure 5: Impact of IFNγ treatment on expression of IFNγ-realted genes.** qPCR analysis of *IFNGR1* and *IFNGR2* relative expression in D18 'UNTR', 'IFNγ', 'As' and 'As + IFNγ' conditions. *n* = 4 biological replicates from 3 control cell lines. No significant differences observed. One-way ANOVA with Tukey's multiple comparison test.

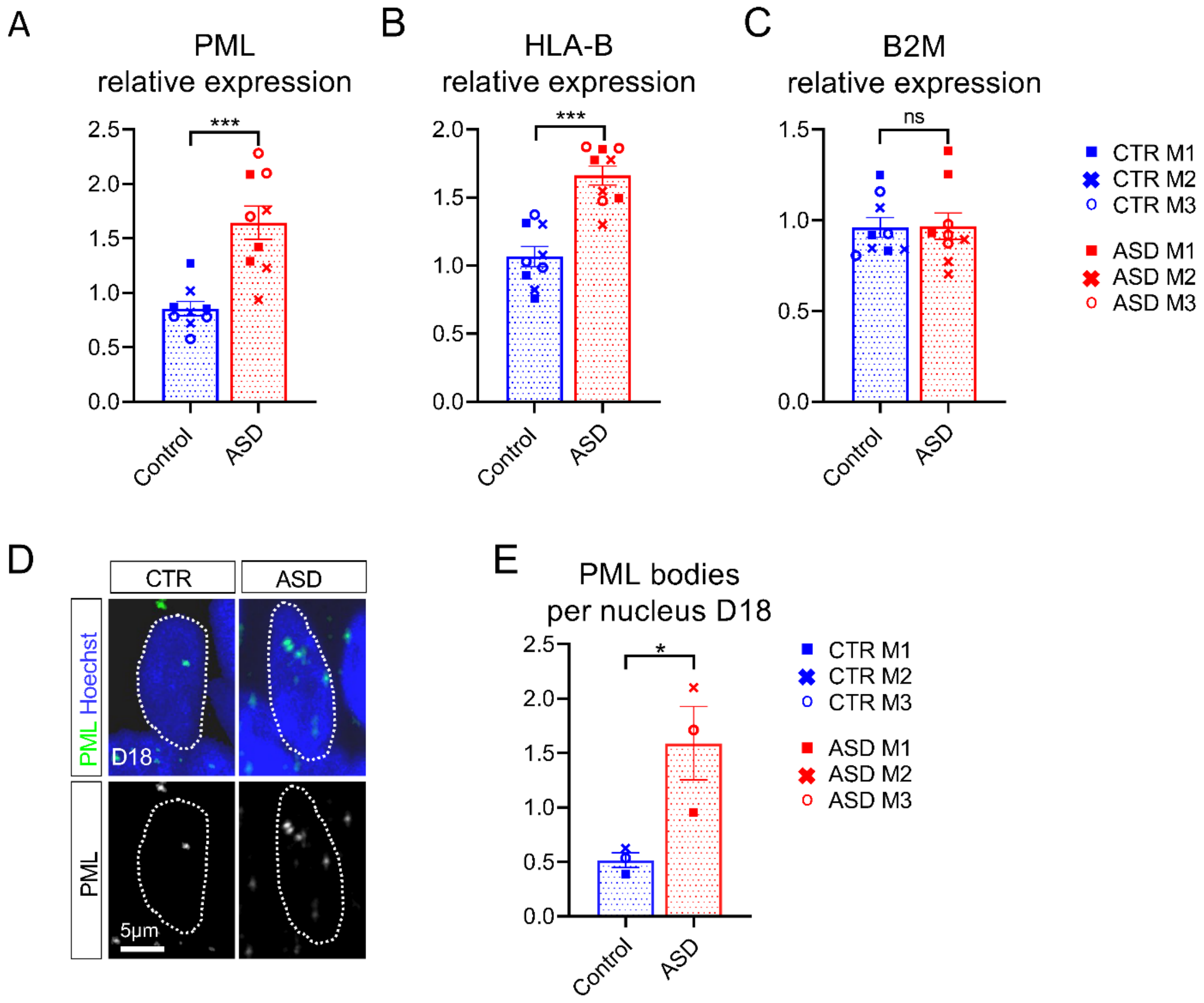


**Supplementary Figure 6: NPCs generated from ASD-hiPSCs display elevated levels of PML and HLA-B expression. (A-C)** QPCR analysis of *PML*, *HLA-B* and *B2M* in day 18 NPCs. For all genes, *n* = 9 biological replicates from 3 control or 3 ASD hiPSC lines. *PML* and *HLA-B* but not *B2M*, showed increased expression in NPCs from ASD hiPSCs. Unpaired t test. **(D)** Confocal images of PML nuclear bodies in D18 NPCs derived from control and ASD hiPSCs. **(E)** Quantification of PML nuclear bodies at D18 in NPCs derived from *n* = 3 control and *n* = 3 ASD hiPSC lines. CTR: 136 cells analysed; ASD: 280 cells analysed. Unpaired t test. Bar chart results are presented as means +/- SEM. **P* < 0.05; ****P* < 0.001; ns: not significant.

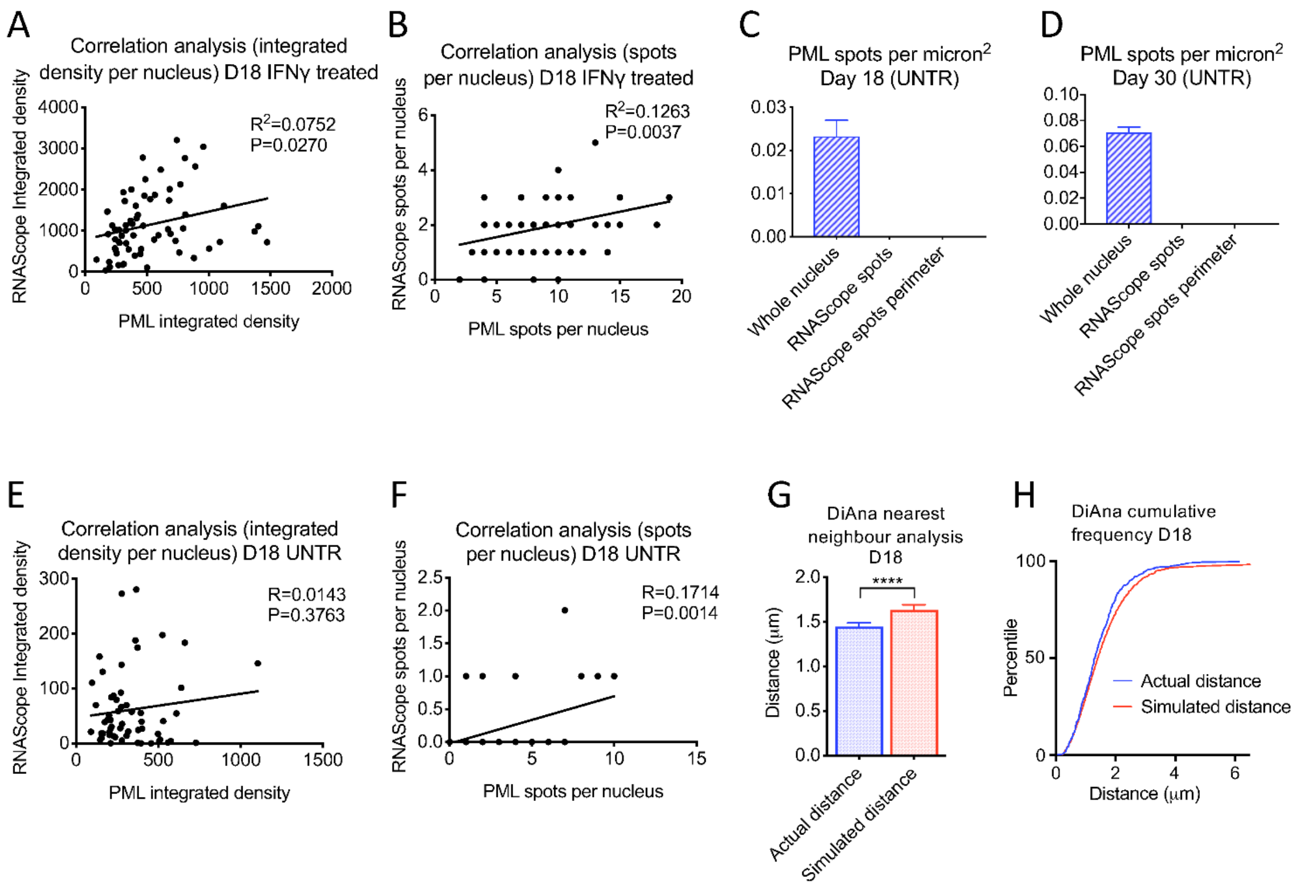


**Supplementary Figure 7: *HLA-B* RNAScope and PML correlation and spatial analysis in D18 UNTR NPCs. (A and B)** Correlation analysis of *HLA-B* RNAScope and PML spots per nucleus and integrated density per nucleus in D18 IFNγ-treated cells. $n$ = 65 cells from 3 control cell lines. Linear regression analysis. **(C and D)** Quantification of PML spots per micron$^2$ in whole nuclei, *HLA-B* RNAScope spots and *HLA-B* RNAScope spot perimeters in D18 NPCs **(C)** and D30 neurons **(D)**, untreated condition. D18 UNTR: $n$ = 59 cells; D30 UNTR: $n$ = 56 cells; 3 control cell lines. **(E and F)** Correlation analysis of *HLA-B* RNAScope and PML integrated density and spots per nucleus in D18 untreated cells. $n$ = 57 cells from 3 control cell lines. Linear regression analysis. **(G and H)** DiAna distance analysis of centre-centre distances from *HLA-B* RNAScope spots to PML spots in real images and following random shuffle of PML and RNAScope spots, using nuclei as a bounding box. n = 405 measurements; 3 control cell lines. Wilcoxon matched-pairs test. Results are presented as means +/- SEM. ****$P$ < 0.0001.
